# TOLLIP resolves lipid-induced EIF2 signaling in alveolar macrophages for durable *Mycobacterium tuberculosis* protection

**DOI:** 10.1101/2020.08.24.263624

**Authors:** Sambasivan Venkatasubramanian, Courtney Plumlee, Kim Dill-McFarland, Gemma L. Pearson, Sara B. Cohen, Anne Lietzke, Amanda Pacheco, Sarah A. Hinderstein, Robyn Emery, Scott A. Soleimanpour, Matthew Altman, Kevin B. Urdahl, Javeed A. Shah

## Abstract

Relative deficiency of TOLLIP expression in monocytes is associated with increased tuberculosis (TB) susceptibility in genetic studies, despite antagonizing host innate immune pathways that control *Mycobacterium tuberculosis* (Mtb) infection. In this study, we investigated the mechanisms by which TOLLIP influences Mtb immunity. *Tollip*^-/-^ mice developed worsened disease, consistent with prior genetic observations, and developed large numbers of foam cells. Selective TOLLIP deletion in alveolar macrophages (AM) was sufficient to induce lipid accumulation and increased Mtb persistence 28 days after infection, despite increased antimicrobial responses. We analyzed sorted, Mtb-infected *Tollip*^-/-^ AM from mixed bone marrow chimeric mice to measure global gene expression 28 days post-infection. We found transcriptional profiles consistent with increased EIF2 signaling. Selective lipid administration to *Tollip*^-/-^ macrophages induced lipid accumulation, and Mtb infection of lipid laden, *Tollip*^-/-^ macrophages induced cellular stress and impaired Mtb control. EIF2 activation induced increased Mtb replication within macrophages, irrespective of TOLLIP expression, and EIF2 kinases were enriched in human caseous granulomas. Our findings define a critical checkpoint for TOLLIP to prevent lipid-induced EIF2 activation and demonstrate an important mechanism for EIF2 signaling to permit Mtb replication within macrophages.

## Introduction

Toll-Interacting Protein (TOLLIP) is a selective autophagy receptor and endosomal sorting protein that was initially identified as a TLR and IL-1R binding protein, (*1*) and chaperones membrane bound receptors and protein aggregates to the autophagosome via endoplasmic reticulum (ER) transport and autophagy (*2, 3*)(*4, 5*). A functionally active single nucleotide polymorphism upstream of the TOLLIP transcriptional start site diminishes TOLLIP gene expression in monocytes and is associated with increased risk for pulmonary and meningeal TB, increased innate immune responses after Mtb infection, and diminished BCG-specific T cell responses in South African infants (*6, 7*). We sought to define the mechanism by which TOLLIP influences Mtb susceptibility, especially during chronic phases of infection. Therefore, we evaluated TOLLIP’ mechanism of effect on pulmonary innate immune responses to Mtb infection and TB severity over time, in small animal models. In these papers, identified a critical checkpoint of lipid homeostasis required for durable control in chronically Mtb-infected macrophages.

The impact of autophagy machinery on Mtb pathogenesis is incompletely understood. After IFNγ activation, autophagy contributes to host defense within murine macrophages by trafficking Mtb to autophagolysosomes for destruction, but this only represents a partial contribution to Mtb clearance (*8, 9*). Autophagy also dampens innate immune responses, including inflammasome and TLR activity (*10*), and the autophagy regulator Atg5 controls neutrophil-induced tissue destruction and immunopathology after Mtb infection (*11, 12*). Autophagy is also induced to clear misfolded proteins and excess lipid as part of the integrated stress response (ISR), which promotes cell survival and is an important mechanism for maintaining cellular function over time, but its role in Mtb pathogenesis is not well understood (*13*). TOLLIP, an autophagy receptor, is associated with TB susceptibility, so understanding its role in macrophage function may provide insight into how autophagy machinery can influence Mtb pathogenesis.

The influence of integrated stress response (ISR) on Mtb pathogenesis is not completely understood. The ISR is induced by excess misfolded protein, lipid, or heat shock, which activate one of four kinases (HRI, GCN2, PERK, and PKR) to phosphorylate EIF2S1 (*14*). EIF2S1 phosphorylation induces dramatic changes in cell translation and function, which may lead to adaptation and cellular survival by inducing autophagy or cell death, if stresses are prolonged or severe (*14, 15*). Multiple lines of evidence suggest that the ISR may prevent durable Mtb control. Granulomas are hypoxic, associated with increased lipid content, and are inflammatory, and each of these phenotypes induces the ISR (*16–18*). The ISR diminishes protein translation and impairs glycolysis, and these phenotypes are associated with decreased macrophage control of Mtb (*19, 20*). Oxidative stress, another inducer of the ISR, in macrophages induces inflammatory overreaction to bacterial products and TNF, which can worsen outcomes from TB infection (*21–24*). Mtb infection of macrophages induces ER stress, but whether this is beneficial by inducing apoptosis or harmful by preventing adequate Mtb control is not certain (*25–27*). In this study, we found that TOLLIP-deficient mice develop selective lipid accumulation in their alveolar macrophages (AM) during Mtb infection, prompting us to test the effects of selective intracellular lipid accumulation on Mtb pathogenesis *in vitro* and *in vivo*. In this absence of clearance, excess lipid induces the ISR and permits Mtb replication, which ultimately leads to Mtb progression.

## Results

### Tollip^-/-^ macrophages develop proinflammatory cytokine bias

In preliminary experiments, we identified a functional single nucleotide polymorphism in the TOLLIP promoter region that was associated with decreased TOLLIP mRNA expression in peripheral blood monocytes and hyperinflammatory cytokine responses after TLR stimulation and Mtb infection (*6, 7, 28*). This variant was also associated with increased risk for pulmonary and meningeal TB in genetic studies (*28*). To link these human genetic observations with our small animal model, we evaluated the functional capacity of *Tollip*^-/-^ macrophages to produce pro- and anti-inflammatory cytokines after TLR stimulation and Mtb infection. We isolated peritoneal macrophages (PEM), plated them *ex vivo* and stimulated them with LPS (TLR4 ligand; 10ng/ml), PAM3 (TLR2/1 ligand; 250ng/ml), or Mtb whole cell lysate (1μg/ml). *Tollip*^-/-^ PEM secreted more TNF than control after all stimulation conditions (**Figure S1A**, LPS p = 0.01, PAM3 p = 0.002, Mtb lysate p = 0.01, n = 9). Conversely, *Tollip*^-/-^ PEM induced less IL-10 than controls (**Figure S1B**, LPS p = 0.01, PAM3 p = 0.02, Mtb lysate p =0.03, n = 9). We infected PEM with live Mtb H37Rv strain (MOI 2.5) overnight and measured TNF, IL-1β and IL-10. After infection, *Tollip*^-/-^ PEM secreted more TNF and IL-1 β, while inducing less IL-10 than controls (MOI 2.5) (**Figure S1C-E**; p=0.03, p=0.03, and p=0.02 respectively), which is consistent with prior human observations (*7, 28*). Thus, murine TOLLIP recapitulated the functional phenotypes of human TOLLIP after Mtb infection in macrophages.

### TOLLIP is required to control Mtb infection

A functionally active variant in the TOLLIP promoter was associated with increased TB susceptibility (*6, 7, 28*). This SNP was also associated with 1) decreased TOLLIP mRNA expression in monocytes, 2) increased innate immune responses to Mtb infection, but 3) diminished BCG-specific CD4+ T cell responses, suggesting multiple possible mechanisms by which it may mediate TB progression (*7*). Therefore, we evaluated how TOLLIP influenced TB outcomes in the knockout mouse model to elucidate the mechanistic underpinnings of TOLLIP in the immune response to Mtb *in vivo*. We infected *Tollip*^-/-^ mice and littermate controls expressing *Tollip*^-/-^ (WT) with 100 cfu of Mtb H37Rv strain via aerosol and monitored weight, bacterial colony forming units (CFU), and survival over time (**Figure 1A**). Lung CFU were diminished two weeks after infection in *Tollip*^-/-^ mice (p < 0.009, n = 5; **Figure 1B**). Four weeks post-infection, Mtb bacterial burden in the lung was diminished in *Tollip*^-/-^ mice compared with controls (**Figure 1C**, n =30 / mouse type; p < 0.020, two-sided t-test). Eight weeks post-infection, this phenotype reversed and *Tollip*^-/-^ mice developed increased bacterial burden (p=0.0011, two-sided t-test) and beyond. By 180 days, lung bacterial burdens were increased ten-fold in *Tollip*^-/-^ mice compared to controls (p = 0.03, t-test; overall difference in groups, p = 0.004, mixed effects model comparing, genotype, time, and TOLLIP expression). Similarly, we found extrapulmonary dissemination of Mtb was delayed in *Tollip*^-/-^ mice, as we were unable to culture Mtb from of 3/5 spleens (**Figure 1D**, p = 0.06, two-sided t-test). Four weeks after infection, bacterial load was decreased in the spleens of *Tollip*^-/-^ mice, but we observed increased bacterial burden in the spleen after eight weeks and onward (**Figure 1E**, p = 0.0092, 2-way ANOVA comparing time and genotype). *Tollip*^-/-^ mice met criteria for euthanasia a median of 228 days after infection (**Figure 1F**; p < 0.0001, n = 10 / group, Mantel-Cox test), whereas WT mice did not lose weight and survived over 250 days after infection. *Tollip*^-/-^ mice lost weight beginning approximately 180 days post-infection as well (**Figure 1G**, p < 0.0001, 2-way ANOVA accounting for genotype and time). Histopathologic analysis of lung sections also revealed differences between early and late immune responses in *Tollip*^-/-^ mice. Although *Tollip*^-/-^ and WT mice developed qualitatively similar inflammatory lesions four weeks post-infection, by eight weeks post-infection, *Tollip*^-/-^ mice displayed increased numbers of cellular structures with a foamy appearance within infiltrates, consistent with lipid-laden “foamy” macrophages (**Figure 1H**). Taken together, *Tollip*^-/-^ mice exhibit early resistance to Mtb infection during the first 2-4 weeks, which might be predicted by the enhanced inflammatory response of *Tollip*^-/-^ macrophages. Paradoxically, during chronic stages of Mtb infection their susceptibility is reversed and *Tollip*^-/-^ mice exhibit higher bacterial burdens than WT controls.

**Figure 1.**
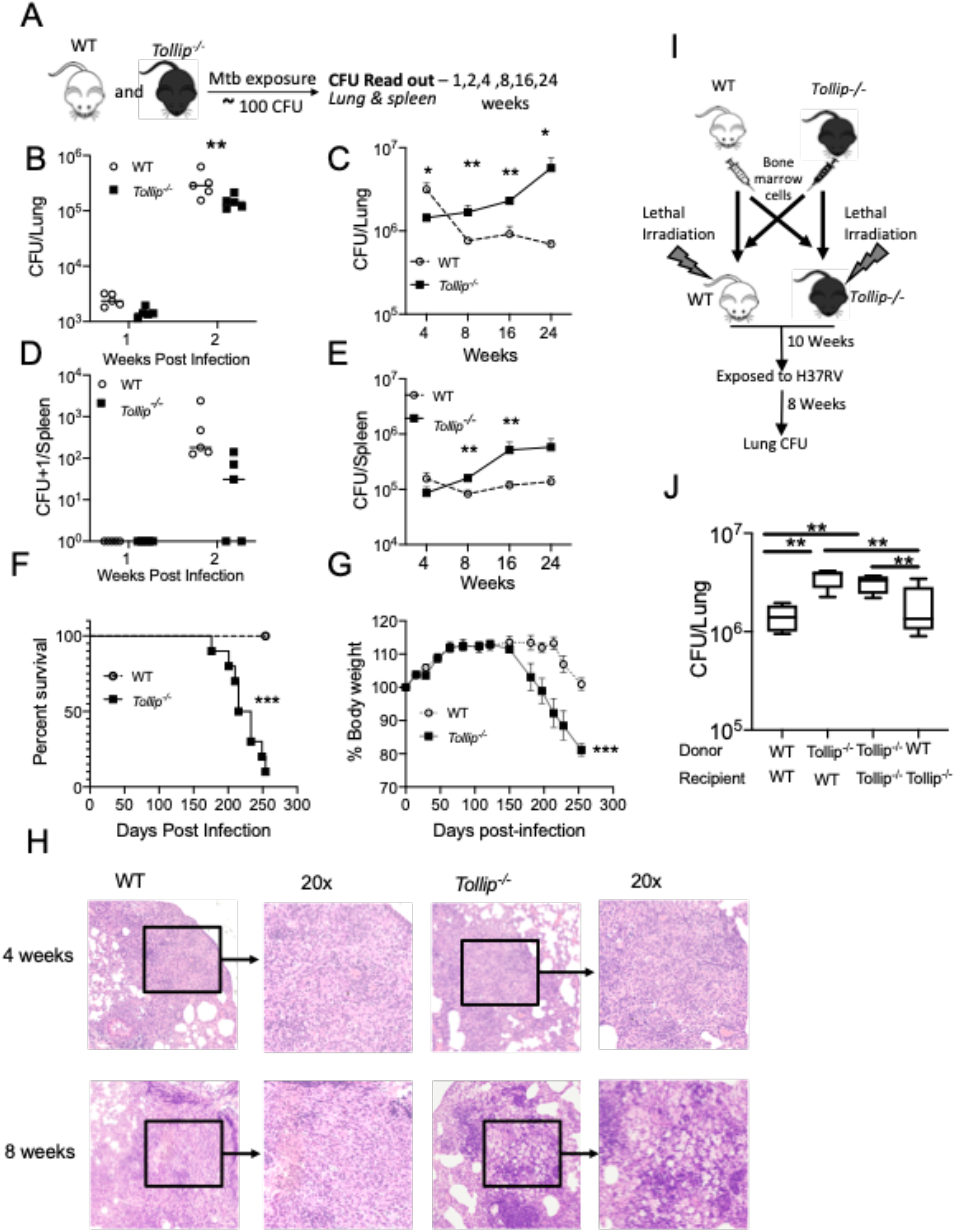
TOLLIP is required for Mtb control in mice and its absence induces foam cell formation. Mice were infected with Mtb H37Rv (50-100 cfu) via aerosol and monitored over time. **A)** Experimental timeline. **B-C)** Lung bacterial burden in WT (*clear circle*) and *Tollip*^-/-^ (*black square*) mice **B)** 1-2 weeks and **C**) 4 weeks and later after infection. Spleen bacterial burden **D**) 1-2 weeks and **E)** 4 weeks and later after infection. * p < 0.05; ** p < 0.01, ***p < 0.001, two-sided t-test at each time point; n = 5 mice/group. Overall associations were determined using a two-sided ANOVA accounting for time and genotype. **F)** Survival curve analysis of WT (*clear circle*) and *Tollip*^-/-^ (*black square*) mice after aerosol infection with 50-100cfu Mtb H37Rv strain. N = 10 mice/group. P < 0.0001, Mantel-Cox test. **G)** Percentage of initial body weight after Mtb aerosol infection *Circle* – WT mice; *square - Tollip*^-/-^ mice. N = 10 / group. **H)** Hematoxylin and eosin staining of Mtb-infected lung tissue from WT and *Tollip*^-/-^ mice 4 and 8 weeks after aerosol infection. **I)** Experimental outline of full bone marrow chimera experiments. **J)** Lung bacterial burden of bone marrow chimeric mice, 8 weeks after aerosol Mtb infection. N = 5 mice/group. * p < 0.05, ** p < 0.01, *** p < 0.001.

Because TOLLIP is ubiquitously expressed (*29*), we generated reciprocal bone-marrow chimeras to assess whether TOLLIP’s expression in hematopoietic or non-hematopoietic cells is required for immunity against Mtb. Lung bacterial burdens were assessed in mice infected with Mtb eight weeks prior (**Figure 1G**). Consistent with our results in non-chimeric mice, *Tollip*^-/-^ bone marrow -> *Tollip*^-/-^ mice exhibited higher lung bacterial burdens that WT -> WT controls. Importantly, *Tollip*^-/-^ -> WT chimeras had elevated bacterial burdens like *Tollip*^-/-^ -> *Tollip*^-/-^ controls, whereas bacterial burdens in WT -> *Tollip*^-/-^ were not significantly different than WT - > WT controls. These data suggest a specific role for TOLLIP in radiosensitive hematopoietic cells during Mtb infection.

### TOLLIP deficiency induces increased intracellular Mtb burden in AM during chronic infection

Having shown that TOLLIP expression is required in hematopoietic cells for Mtb immunity, we next sought to identify the specific hematopoietic cell types influenced by TOLLIP expression. In prior studies, we showed that TOLLIP deficiency promotes diminished Mtb replication within macrophages *in vitro*, which correlated with its role in diminishing proinflammatory cytokine responses (*6, 30*). However, given the opposing results in both human genetic studies and mouse infectious challenge, we investigated whether TOLLIP was required for Mtb control in distinct pulmonary macrophages within the lung over the disease course. We infected *Tollip*^-/-^ and WT mice with Mtb expressing an mCherry fluorescent reporter (50-100 CFU) via aerosol and four and eight weeks after infection, we measured lung myeloid cell populations and the proportion of cells infected with Mtb using an adaptation of recently devised gating strategies (**Figure S2,** (*31, 32*)). We found no difference in the proportion of AM (CD11c+SiglecF+), monocyte-derived macrophages (MDM; SiglecF-CD11b+CD11c+MHCII+), interstitial macrophages (IM; SiglecF-CD11b+CD11c-MHCII+), neutrophils (PMN; SiglecF-CD11b+Ly6G+), or dendritic cells (DC; SiglecF-CD11b-CD11c+MHCII+) found in the lung between WT and *Tollip*^-/-^ mice during the course of Mtb infection (**Figure 2A-B**). When we assessed the infected cells, however, we observed a lower percentage of mCherry+ AM and PMN infected with Mtb in *Tollip*^-/-^ mice, compared to WT mice four weeks post-infection (**Figure 2C**, p = 0.01 and p = 0.02, respectively; n =5). By eight weeks, however, there was a reversal and a significantly greater proportion of Mtb-infected AM and MDM from *Tollip*^-/-^ mice (**Figure 2D**, p = 0.03 and p = 0.01, respectively). Thus, although the overall proportion of myeloid populations were similar in WT and *Tollip*^-/-^ mice, some myeloid cell subtypes, including AM, were relatively less infected at early timepoints, and relatively more infected at later timepoints.

**Figure 2.**
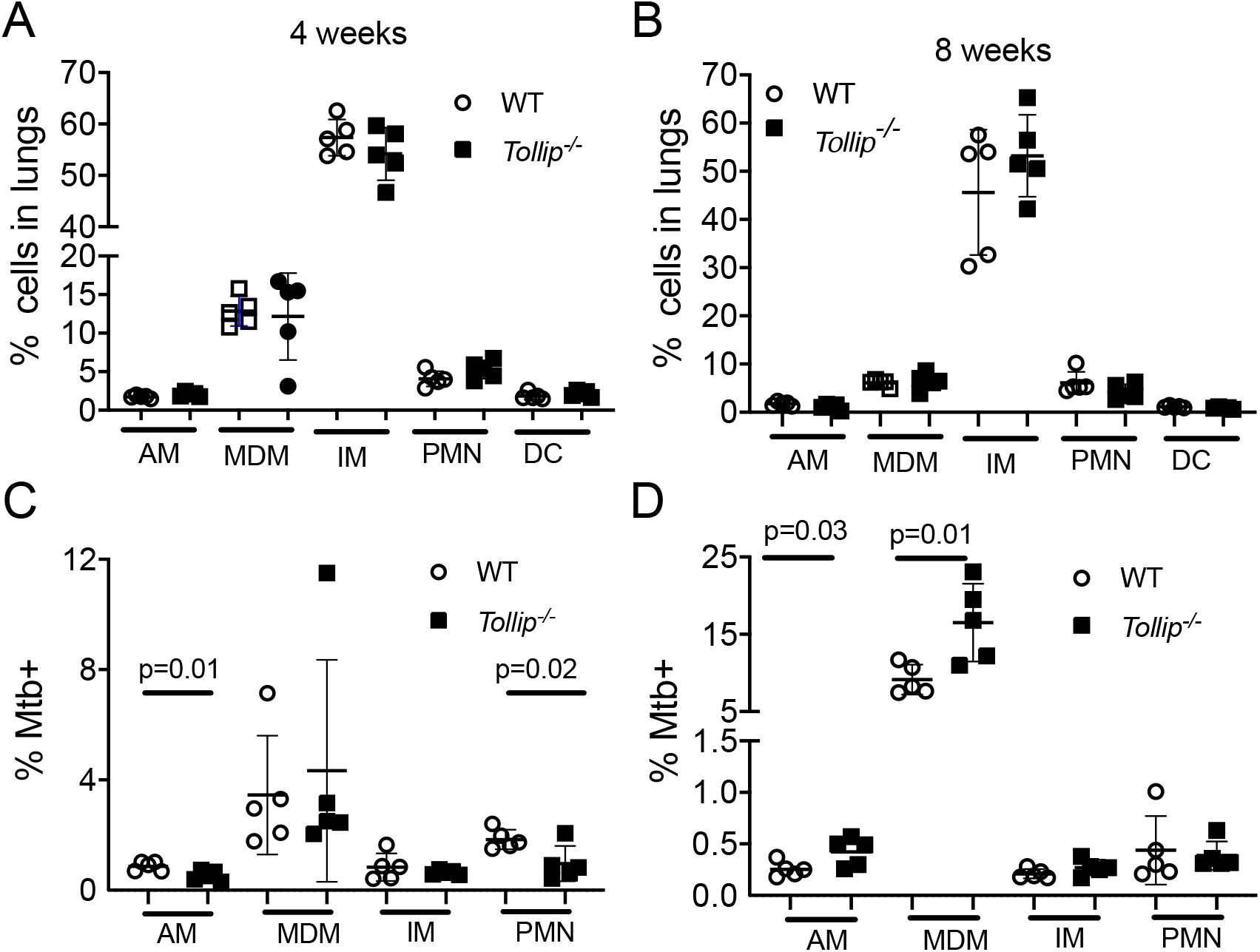
Mtb preferentially infects *Tollip*^-/-^ myeloid cells 8 weeks after aerosol infection. Lungs from Mtb-infected mice were gently dissociated, and flow cytometry was performed. Gating strategy described in **Figure S2**. The proportion of lung resident myeloid cells from WT (*clear circle*) and *Tollip*^-/-^ (*black square*) mice was compared **A)** 4 and **B)** 8 weeks after aerosol infection with 50-100 cfu mCherry-expressing Mtb. The proportion of each cell subset infected with Mtb (mCherry+) was compared between groups **C)** 4 and **D)** 8 weeks post infection. N = 5 mice/group. Experiment was performed three times independently. Statistical significance determined using Student’s t-test. AM – alveolar macrophage, MDM – monocyte-derived macrophage, IM – interstitial macrophage, PMN – neutrophil, DC – dendritic cell.

### TOLLIP deficiency induces and maintains increased NOS2 expression in lung-resident myeloid cells after Mtb infection

Next, we compared the functional capacity of lung-resident myeloid cells in WT and *Tollip*^-/-^ mice. Four weeks post-infection, Mtb-infected AMs from *Tollip*^-/-^ mice demonstrated significantly increased NOS2 median fluorescence intensity when compared to WT littermates (**Figure 3A**, p = 0.01, n = 5), and we observed a trend toward increased NOS2 from Mtb infected PMNs in *Tollip*^-/-^ mice. Eight weeks after infection, NOS2 expression was similar in all myeloid cell types across WT and *Tollip*^-/-^ mice (**Figure 3B**). We measured expression of MHC Class II and CD80 between Mtb-infected and uninfected myeloid cells and found that, while MHC Class II expression from each myeloid cell type was similar (data not shown), CD80 expression was significantly diminished on Mtb-infected *Tollip*^-/-^ AM (p = 0.008), MDM (p = 0.05), and PMN (p =0.05) four weeks post-infection (**Figure 3C**, n = 5/group) and in *Tollip*^-/-^ Mtb-infected AM and MDM (**Figure 3D,** p=0.002 and p=0.004, respectively) eight weeks post-infection. In summary, NOS2 expression, a marker associated with antimicrobial activity, was increased at early timepoints in Mtb-infected AM, but this was not sustained at later timepoints. In contrast, CD80 expression, a molecule critical for T cell costimulation, was diminished at both early and late timepoints in multiple Mtb-infected myeloid cell populations in *Tollip*^-/-^ mice.

**Figure 3.**
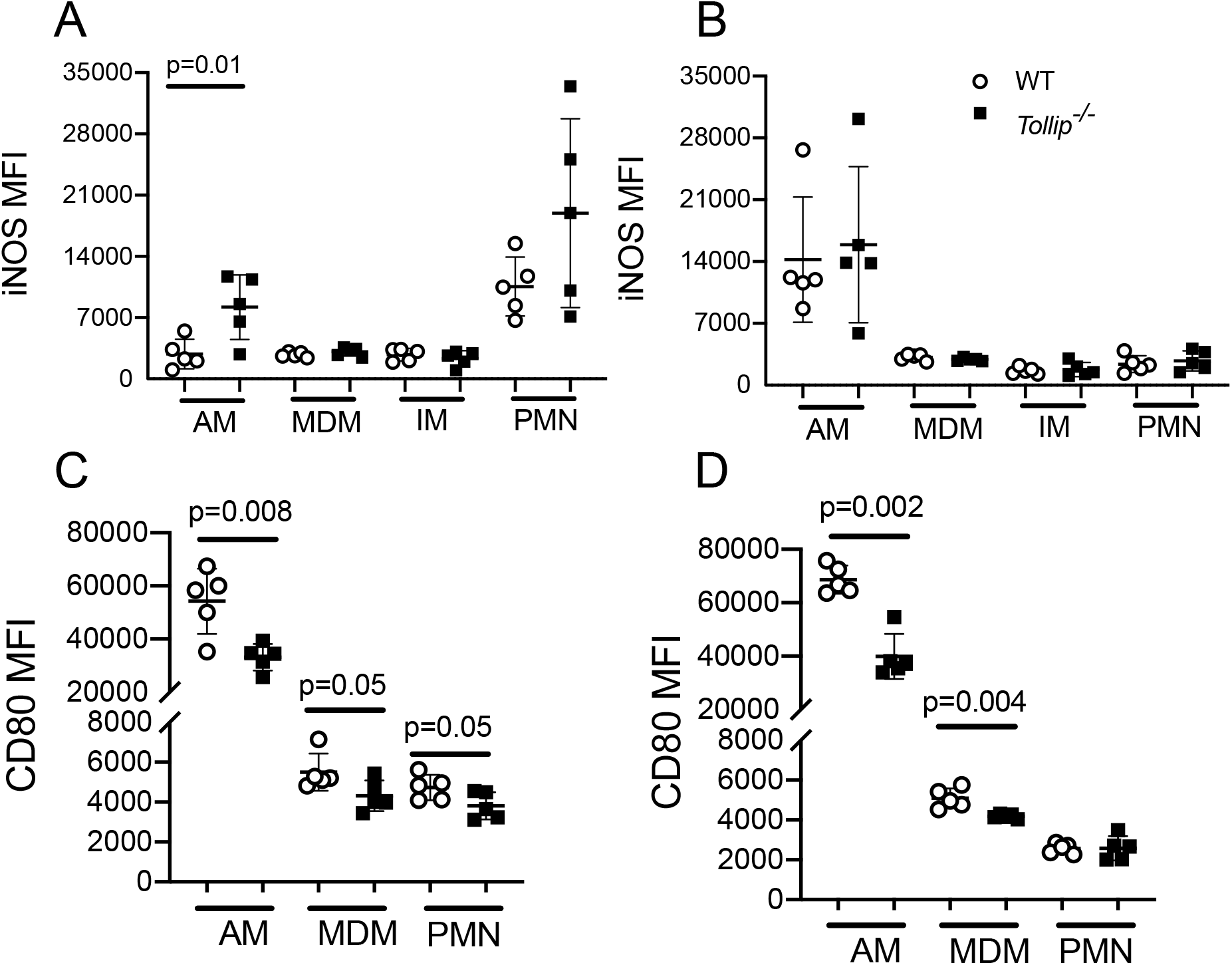
*Tollip*^-/-^ lung-resident myeloid cells develop enhanced antimicrobial responses after Mtb infection. Please see **Supplemental Figure 2** for gating strategy to identify myeloid cell subsets. **A and B)** Intracellular NOS2 protein expression was measured in Mtb-infected lung-resident myeloid cells (mCherry+) by flow cytometry **A)** 4 and **B)** 8 weeks post infection. CD80 expression was measured **C)** 4 and **D)** 8 weeks post infection. Median fluorescence intensity was compared using a two-sided t-test. N = 5 mice/group. WT (*clear circle*); *Tollip*^-/-^ (*black square*). Experiment was repeated three times independently. AM – alveolar macrophage, MDM – monocyte-derived macrophage, IM – interstitial macrophage, PMN – neutrophil, DC – dendritic cell.

### Intrinsic expression of TOLLIP in AM regulates immunity against Mtb

Our findings in Mtb-infected *Tollip*^-/-^ mice suggested that TOLLIP expression in AM may regulate their ability to control Mtb. However, in global *Tollip*^-/-^ knockout mouse models, these results could also reflect indirect effects of TOLLIP-deficiency in other cell types, or bacterial burden differences between WT and *Tollip*^-/-^ mice. To directly assess the intrinsic role of TOLLIP in distinct myeloid populations, we generated mixed bone marrow chimeric mice by combining bone marrow from F1 generation *WT* mice (CD45.1+CD45.2+) in a 1:1 ratio with *Tollip*^-/-^ (CD45.2+) bone marrow and transplanting this mix into CD45.1 + recipient mice (**Figure 4A**). After allowing 10 weeks for immune reconstitution and confirmation of equal proportions of immune cells from each lineage (**Figure 4B**), we infected these mice with 50-100cfu Mtb-mCherry and evaluated the induction and maintenance of the innate immune response to Mtb. Our flow gating strategy is demonstrated in **Figure S3**. We recapitulated the pattern of early innate immune responses to Mtb (*32*). 14 days after infection, AM were the primary cell infected, but increasing proportions of PMN and MDM became infected over time, as described in prior publications (**Figure 4C;** (*32*). We measured the proportion of *WT* and *Tollip*^-/-^ myeloid cells infected with Mtb over time. 14 days post infection, a greater percentage of *WT* AM were infected with Mtb compared with *Tollip*^-/-^ AM (**Figure 4D**, p = 0.001; paired t-test, n = 5, representative of 3 independent experiments). In contrast, the percentage of Mtb-infected WT and *Tollip*^-/-^ MDM and PMN were similar. By d28, however, we observed a reversal of the AM phenotype; a greater percentage of *Tollip*^-/-^ AM were infected compared to their WT counterparts (**Figure 4E**, p = 0.015). Further, despite the fact that AM make up a relatively modest percentage of cells infected by Mtb, these cells are preferentially infected, suggesting that they remain a preferential replicative niche throughout Mtb infection. A modest increase in the proportion of Mtb-infected *Tollip*^-/-^ PMN was also observed at d28 (**Figure 4E**, p = 0.02). Consistent with our findings in global *Tollip*^-/-^ mice, we found that NOS2 expression was increased in *Tollip*^-/-^ AM in the mixed chimeras as compared to WT AM (**Figure 4F**, p = 0.04; paired t-test; n=5). Overall, these findings show a critical AM-intrinsic role for TOLLIP in the regulation of their immune response to Mtb-infection and that the nature of this role changes over time. At early timepoints, TOLLIP deficiency in AM restricts Mtb accumulation, however, at later timepoints this is reversed and TOLLIP-deficiency in AM promotes cell-intrinsic Mtb infection.

**Figure 4.**
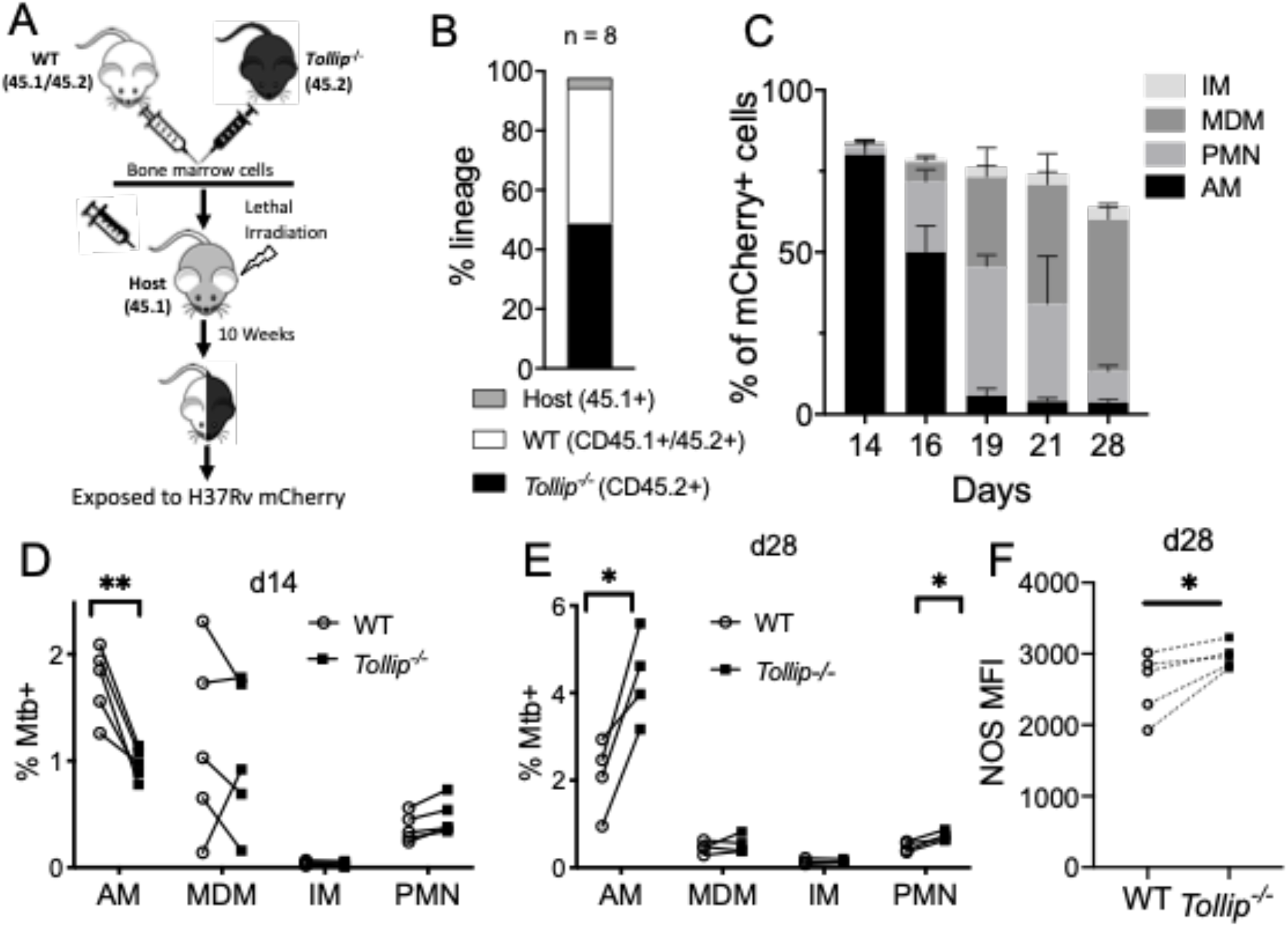
*Tollip*^-/-^ AM become preferentially infected with Mtb over time. *WT:Tollip*^-/-^ mixed bone marrow chimeric mice were infected with Mtb expressing mCherry (50-100 cfu) and followed. Gating strategy can be found in **Figure S3**. **A)** Experimental strategy and timeline. Mixed bone marrow chimeras were generated by mixing 1:1 ratios of WT (CD45.1+/CD45.2+) and *Tollip*^-/-^ (CD45.2+) mice and transferred to a CD45.1 + murine host. **B)** Frequency distribution of CD45 expression in naïve mixed bone marrow chimeric mice. **C)** Composition of mCherry+ lung leukocytes at the indicated time point. **D and E)** Distribution of mCherry+ cells at **D)** d14 and **E)** d28 post-infection in mixed bone marrow chimeric mice. **F)** Median fluorescence intensity of nitric oxide synthase (NOS) in AM at d28 post-infection from mixed bone marrow chimeric mice. N = 4 or 5 mice for each time point as indicated; experiments were conducted three times independently. * p < 0.05, ** p < 0.005. Significance determined by paired two-sided t-test. AM – alveolar macrophage, MDM – monocyte-derived macrophage, IM – interstitial macrophage, PMN – neutrophil.

### Tollip^-/-^ *AM develop increased EIF2 signaling responses after Mtb infection*

The early resistance of *Tollip*^-/-^ AM to Mtb infection can be explained by the observation that *Tollip*^-/-^ macrophages exhibit elevated inflammatory responses and effector functions to Mtb infection, but the reason Mtb persists within AM at later timepoints is less clear. Thus, we evaluated the transcriptional networks associated with loss of Mtb control to identify alternate TOLLIP mechanisms that alter AM function. We sorted infected and uninfected *Tollip*^-/-^ and WT AM from mixed bone marrow chimeric mice at d28 post-infection for analysis by RNA-Seq (sorting strategy is shown in **Figure S4**). We identified 194 differentially expressed genes (DEG) between WT and *Tollip*^-/-^ Mtb-infected AM (**Figure 5A**; FDR < 0.05; gene names at **Table S2**,) and 157 DEGs between WT and *Tollip*^-/-^ Mtb-uninfected “bystander” AM using the same stringent FDR criteria. (**Figure 5B;** list in **Table S2**). The full gene list is available at https://github.com/altman-lab/JS20.01. We investigated genotype-specific gene sets in each group by applying weighted gene coexpression network analysis (WGCNA) and comparing Mtb-infected or -uninfected “bystander” AM by TOLLIP genotype. We used supervised-WGCNA, first identifying and subsetting to 3899 DEGs that met a more lenient FDR cutoff < 0.3 comparing among the 4 groups, to identify 17 distinct coexpression modules containing 54-583 genes per module (**Table S3**). The average expression of genes in each module was then modeled comparing the two AM subsets (WT vs *Tollip*^-/-^) in each condition (infected or uninfected; **Figure 5C**). To understand the biological function of the genes within each module, we tested for enrichment of genes in the curated Broad Institute MSigDB Hallmark gene sets, which represent well-defined biological states and processes that display coherent expression (*33*). Multiple pathways were enriched in each module, as visualized by percent of genes in each module that map to Hallmark terms significant for at least one model (FDR < 0.05; **Figure 5D, Table S4**). Inflammatory/immune pathways (IFNγ response, TNFα signaling via NF-kB, inflammatory response, and IL2/STAT5 signaling) were enriched in modules with decreased expression in *Tollip*^-/-^ AM, consistent with the prior observation that TOLLIP increases inflammation due to post-translational trafficking of immune receptors. Thus, decreased transcript here is consistent with transcriptional feedback loops to this activity and expected, based on experimental data regarding TOLLIP’s mechanism of action (*1*). By contrast, the unfolded protein response, adipogenesis, mTORC1 signaling, oxidative phosphorylation, and fatty acid metabolism were enriched in modules with increased expression in *Tollip*^-/-^ AM.

**Figure 5.**
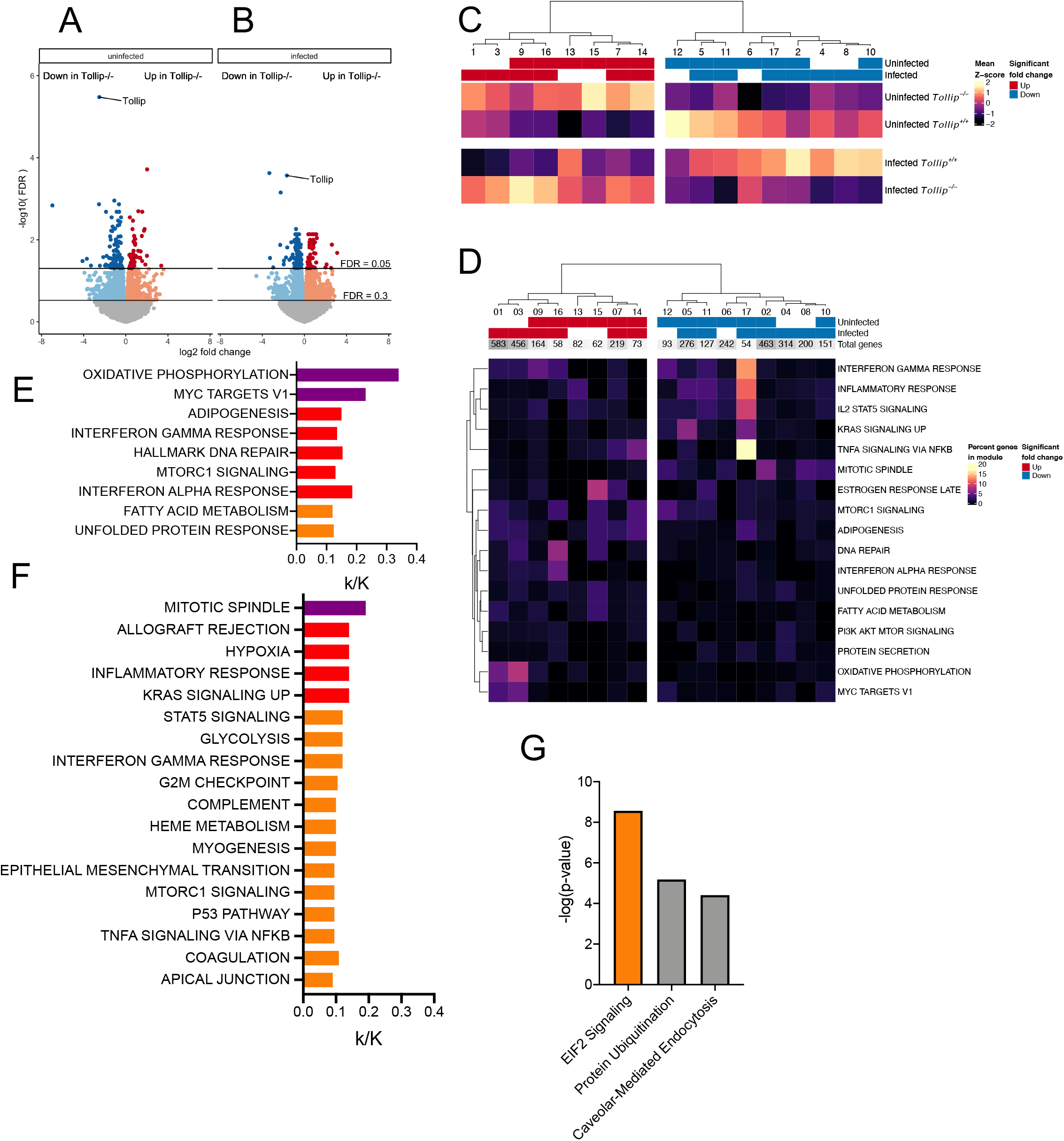
Global gene expression analysis of Mtb-infected *Tollip*^-/-^ AM 28 days after infection demonstrates increased EIF2 signaling. Mixed bone marrow chimeric mice were infected with mCherry-expressing Mtb and 28 days after infection, Mtb-infected and Mtb-uninfected WT and *Tollip*^-/-^ AM were sorted and RNA-seq was performed. Sorting strategy can be found in **Figure S3**. **A)** Volcano plot of gene expression (log2 fold change) and significance (-log10(FDR)) for genes between *Tollip*^-/-^ (CD45.2) and WT (CD45.1+CD45.2+), Mtb-uninfected and **B)** Mtb-infected AMs. Horizontal lines indicate significant genes (FDR < 0.05) and genes used in WGCNA modules (FDR < 0.3) **C)** Heatmap of mean module z-score for infection: TOLLIP groups. Red bars indicate increased expression in *Tollip*^-/-^ AM and blue bars indicated decreased expression in *Tollip*^-/-^ AM, separated by infection status. Modules (columns) were average hierarchical clustered. **D)** Heat map of enrichment of significant Hallmark gene sets (FDR q-value < 0.05) in modules. Color indicates percent of genes in a module that mapped to a given Hallmark. The total number of genes in each module is indicated above. Modules (columns) were clustered as in **C)** and Hallmark terms (rows) were average hierarchical clustered based on enrichment percentages. Red and blue bars are as in **C). E-F)** Bar graph of the Hallmark terms enriched in modules with **E)** increased expression in Mtb-infected, *Tollip*^-/-^ AM or **F)** decreased expression in both Mtb-infected and bystander AM. Bar length indicates the proportional enrichment of each pathway. Color indicates degree of significance: *purple* – FDR q < 10^-16^; *red* -- q < 10^-8^; *orange* – q < 10^-4^. **G)** Ingenuity Pathway Analysis of upstream regulatory pathways enriched by data for Mtb-infected cells. *Orange* bar indicates increased expression in *Tollip*^-/-^ AM. *Gray* bars indicate no directional data available.

To summarize the functional interpretations of RNA sequencing results, we merged modules into 6 subgroups according to a specific pattern of either significantly increased or decreased expression in *Tollip*^-/-^ vs WT AM from 1) Mtb-infected AM only, 2) Mtb-uninfected bystander AM only, or 3) both Mtb-infected and -uninfected populations. We similarly evaluated for pathway enrichment in Hallmark gene sets. The modules showing increased gene expression in *Tollip*^-/-^ Mtb-infected cells were significantly enriched with oxidative phosphorylation (FDR q = 3.96×10^-56^), adipogenesis (q = 2.46×10^-13^), MTORC1 signaling (q = 1.95×10^-10^), fatty acid metabolism (q = 2.52×10^-7^), and unfolded protein response (q = 9.33×10^-6^; **Figure 5E**). The modules showing decreased expression from both Mtb-infected and bystander populations were significantly enriched for hypoxia (q = 1.93×10^-11^) and glycolysis (q = 1.12×10^-8^), mitotic spindle (q = 1.93×10^-19^) and G2M checkpoint (q = 1.03×10^-6^), both cell cycle transcriptional programs, and allograft rejection (q = 1.93×10^-11^), inflammatory response (q = 1.93×10^-11^), IL-2/STAT5 signaling (q = 1.12×10^-8^), IFN-γ response (q = 1.12×10^-8^; **Figure 5F**), which are consistent with multiple immune and inflammatory signaling pathways. The other groups did not demonstrate any significant enrichment for Hallmark gene sets. We performed Ingenuity Causal Network Analysis on genes enriched in Mtb-infected AM (FDR q < 0.05) to identify the upstream regulatory pathways responsible for the observed effects in Mtb-infected *Tollip*^-/-^ AM. We found that the strongest association with increased EIF2 Signaling pathway (**Figure 5G**; z-score 2.714, p = 2.71×10^-9^), EIF2 complex signaling as the primary responsible upstream pathway for phenotypes observed in **Figure 5E**, among Mtb-infected AM.

### *Lipid accumulation in* Tollip^-/-^ *AM permits intracellular Mtb replication*

Bioinformatic analysis suggests that EIF2 signaling, a master switch for the cellular stress response, is the upstream pathway with the strongest association with *Tollip*^-/-^ Mtb-infected AM during chronic Mtb infection. Given the observation of increased foam cells in *Tollip*^-/-^ mice after Mtb infection, we hypothesized that macrophages lacking TOLLIP accumulated excess lipid, which induced EIF2, followed by and maladaptive responses to Mtb. During lipid excess, autophagy and proteosomal activity are induced to control lipid droplet (LD) accumulation (*34*). We first tested the capacity of *Tollip*^-/-^ macrophages to perform bulk macroautophagy in bone marrow derived macrophages (BMDMs) of WT or *Tollip*^-/-^ mice by measuring protein levels of LC3II and p62. To evaluate flux through autophagy, we treated BMDMs with the vATPase inhibitor Bafilomycin A (BafA), which prevents autophagosome/lysosome fusion. As expected, levels of the autophagosome protein marker LC3II were significantly upregulated following BafA treatment. Notably, *Tollip* deletion did not significantly affect levels of LC3II, nor the induction of LC3II by BafA treatment (**Figures 6A-B**). We also assessed levels of the adaptor protein p62, which both targets cargo to autophagosomes for clearance and is itself cleared during autophagy. We again observed similar levels of p62 in both groups, without significant differences in the rise of p62 following BafA treatment (**Figures 6B-C**). However, we did note a non-significant trend towards increased BafA-induced p62 accumulation in *Tollip*^-/-^ cells that could reflect compensation for Tollip deficiency given analogous roles for the proteins (*2, 30*). Nonetheless, these studies indicate that TOLLIP is dispensable for macroautophagy and autophagic flux in macrophages. We also evaluated the impact of TOLLIP on selective autophagy of Mtb within macrophages. We infected TOLLIP-deficient and control macrophage cell lines, regularly used in our lab (*6*), with Mtb expressing mCherry and found no differences between these groups in the proportion of Mtb colocalizing to LC3+ autophagosomes (**Figure S5**).

**Figure 6.**
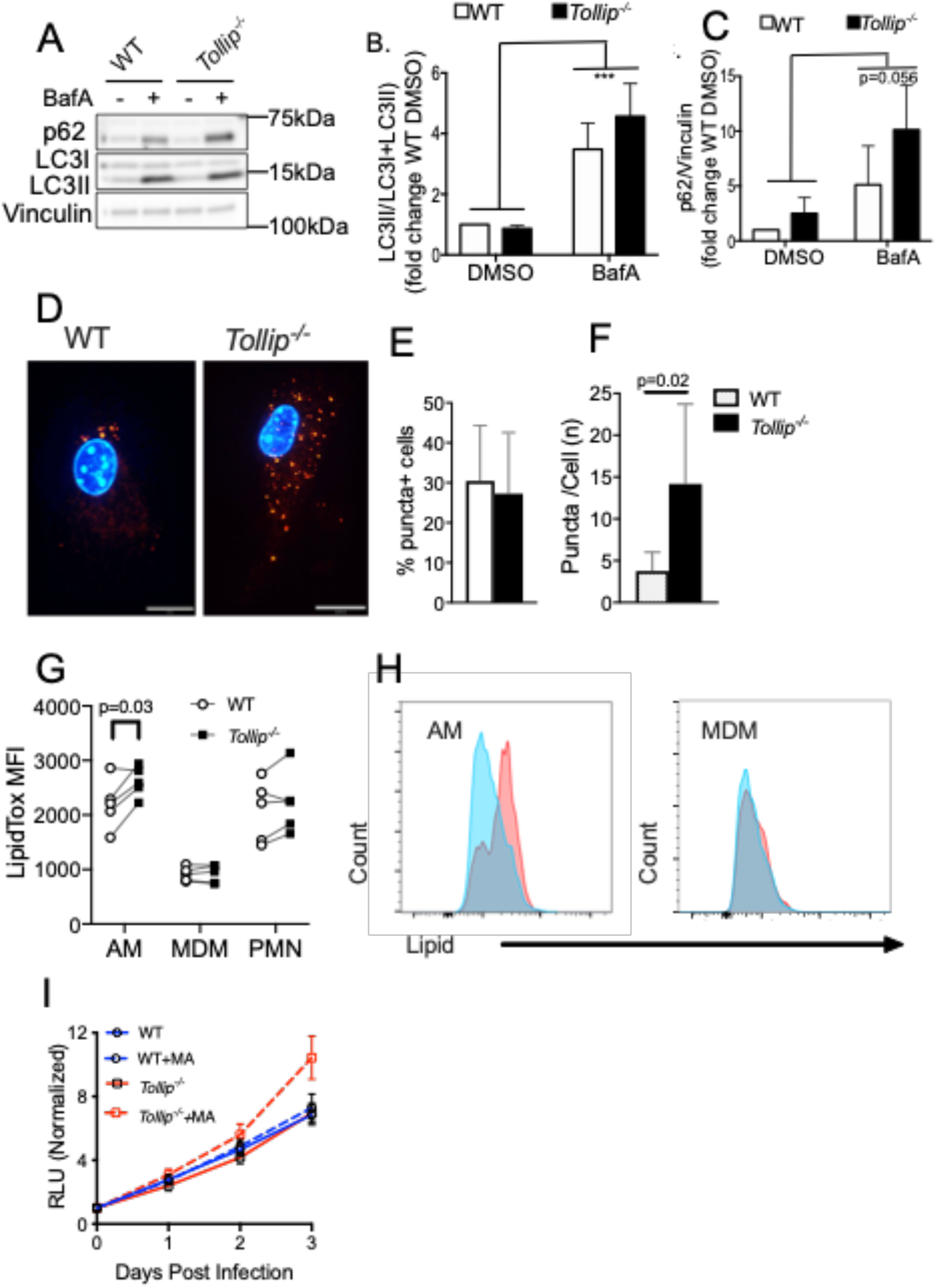
*Tollip*^-/-^ AM accumulate excess lipid that creates a permissive environment for Mtb replication. Bone marrow from WT (*black bars*) or *Tollip*^-/-^ (*white bars*) were differentiated *ex vivo* to macrophages using 40ng/mL mCSF for 7-9 days. Following differentiation, BMDMs were treated for 6hr with or without 250nM Bafilomycin A (BafA). **A**) Representative western blot results showing protein levels of LC3I/II and p62. **B**) Quantification of protein levels (by densitometry) of LC3II using total LC3 (I+II) as a loading control. ***p<0.001 2-way ANOVA for an effect of BafA **C**) Quantification of p62 levels (by densitometry) using vinculin as a loading control. p=0.056 2-way ANOVA effect of BafA. **D-F**) WT and *Tollip*^-/-^ peritoneal exudate macrophages (PEM) were isolated and incubated with mixed mycolic acids (MA) for 72hrs. PEM were stained with LipidTOX Red dye (*red*) and DAPI (*blue*), fixed, and visualized with fluorescent microscopy. **D)** Representative image for of lipid droplet (LD) accumulation in WT and *Tollip*^-/-^ PEM. **E)** Percentage of cells with visible LD, of 100 cells imaged. Bar – error bars (mean ± SD). **F)** Total number of LD per cell, of 100 cells with detectable LD. Bar – error bars (mean ± SD). **G)** Median fluorescence intensity of alveolar macrophages (AM) and monocyte-derived macrophages (MDM) stained with LipidTox neutral lipid stain from mixed bone marrow chimeric mice 28 days after Mtb infection. Infection and gating were performed as in **Figure 5**. Lines connect WT and *Tollip*^-/-^ AM from the same mouse. Significance calculated using two-sided paired t-test. **H)** Representative histogram of AM and MDM lipid staining from one experiment (WT – blue; *Tollip*^-/-^ – red). **I** Luminescence over time in WT and *Tollip*^-/-^ PEM’s incubated with exogenous lipid - MA, mycolic acid (10μg/ml) and infected with Mtb H37Rv expressing *lux* luminescence gene (MOI 1). PEMs were isolated and incubated with MA for 72 hr before infection. Intracellular replication was measured by luminescence for each group over the next three days in quadruplicate and repeated three times to assess reproducibility. Statistical significance was determined by 3-way ANOVA accounting for time, genotype, and lipid administration.

We then evaluated the capacity for TOLLIP to promote lipid clearance overall. We incubated WT and *Tollip*^-/-^ PEM with the mycobacterial cell wall product mycolic acid (MA; 10mg/ml) for 72 hours, stained with fluorescent dye against neutral lipid, and measured the number of LD per cell by microscopy. *Tollip*^-/-^ PEM displayed significantly increased numbers of LD/cell (**Figure 6D**). We noted no difference in the proportion of PEM with quantifiable LD, indicating equal lipid uptake between cells (**Figure 6E**), but a significant increase in the number of LD in *Tollip*^-/-^ PEM (**Figure 6F**, p = 0.02). Next, we evaluated if lung macrophages demonstrated this effect as well. We stained single cell suspensions from the lungs of mixed bone marrow chimeric mice at d28 post-infection for neutral lipid, using gating described in **Figure S4**. We found that *Tollip*^-/-^ AM accumulated significantly more lipid *in vivo* than WT AM, irrespective of their infection with Mtb, while MDM did not significantly accumulate intracellular lipid (**Figure 6G**, p = 0.03; n = 5). A representative histogram for lipid accumulation in AM and MDM is displayed (**Figure 6H**; WT AM – *blue*; *Tollip*^-/-^ AM – *red*).

We measured the impact of lipid accumulation on stress signal transduction and intracellular Mtb replication. First, we confirmed that MA is associated with increased stress responses from BMDM after Mtb infection. In both WT and *Tollip*^-/-^ BMDM, canonical stress signal transduction transcript expression is stable or decreased 24hr after Mtb infection. In WT BMDM, the addition of excess MA does not influence this expression and *Tollip*^-/-^ BMDM demonstrate decreases in expression after infection. However, after exposure to excess MA, followed by overnight live Mtb infection, *Ern1* (**Figure S6A**, p = 0.0007), *Eif2ak3* (**Figure S6B**, p = 0.004), and *Atf6* expression (**Figure S6C**, p = 0.04) is significantly and selectively increased in *Tollip*^-/-^ cells. (**Figure S6A-C**) We infected WT and *Tollip*^-/-^ PEM with a luminescent strain of Mtb H37Rv (Mtb-*lux*, MOI 1; gift of Jeffrey Cox), incubated with or without MA or BSA-conjugated peptidoglycan (PG-BSA) as a non-lipid control. *Tollip*^-/-^ PEM incubated with MA (**Figure 6I**, p = 0.023-way ANOVA accounting for TOLLIP and MA) demonstrated increased Mtb replication compared with *Tollip*^-/-^ PEM in the absence of excess lipid or lipid-incubated WT PEM. Taken together, these results show that TOLLIP prevents lipid accumulation within AM, which induces ER stress, and that dysregulation of the ER stress during chronic infection promotes intracellular Mtb persistence and replication.

### EIF2 Signaling Permits Mtb Replication in Macrophages

After determining that TOLLIP permitted lipid accumulation, which induced cellular stress, we next evaluated the impact of EIF2 activation directly on intracellular Mtb replication. EIF2 is required for protein translation and either complete knockout or activation of this pathway is lethal (*35*). Therefore, we measured effects of this pathway with raphin-1 treatment, a novel inhibitor of protein phosphatase-1, that effectively enhances EIF2 phosphorylation and activation without inducing cell death (*36*). Raphin-1 treatment had no effect on cell death in uninfected macrophages, nor did it influence Mtb replication in broth culture (data not shown). We treated WT PEM with raphin-1 (10μM) or control, then infected these cells with Mtb-*lux* (MOI 1) and followed luminescence over time. Raphin-1 treatment increased Mtb replication in WT PEM (**Figure 7A**; p = 0.0002, 2-way ANOVA accounting for time and treatment). As demonstrated previously, MA alone did not increase Mtb replication in WT PEM, but raphin-1 treatment increased Mtb replication, without evidence of any additive effect with cotreatment with MA and raphin-1 together (**Figure 7B**). Raphin-1 treatment of *Tollip*^-/-^ PEM was associated with increased Mtb replication in *Tollip*^-/-^ PEM compared with WT PEM (**Figure 7C**, p = 0.0475, 3-way ANOVA accounting for TOLLIP, raphin-1, and time). We treated *Tollip*^-/-^ PEM with MA, which was associated with increased impaired Mtb luminescence compared with PEM 72 hr after infection (**Figure 7D**; p = 0.0147, two-sided t-test). Raphin-1 treatment of *Tollip*^-/-^ PEM induced further increases in Mtb replication above MA (**Figure 7D**, p = 0.12). MA and raphin-1 treatment together in *Tollip*^-/-^ PEM induced no synergistic increases in replication, compared with MA alone, after 72 hr (**Figure 7D**). These data demonstrate that EIF2 signaling permits increased Mtb growth, and that decreased TOLLIP expression increases the potency of raphin-1 toward Mtb replication within macrophages.

**Figure 7.**
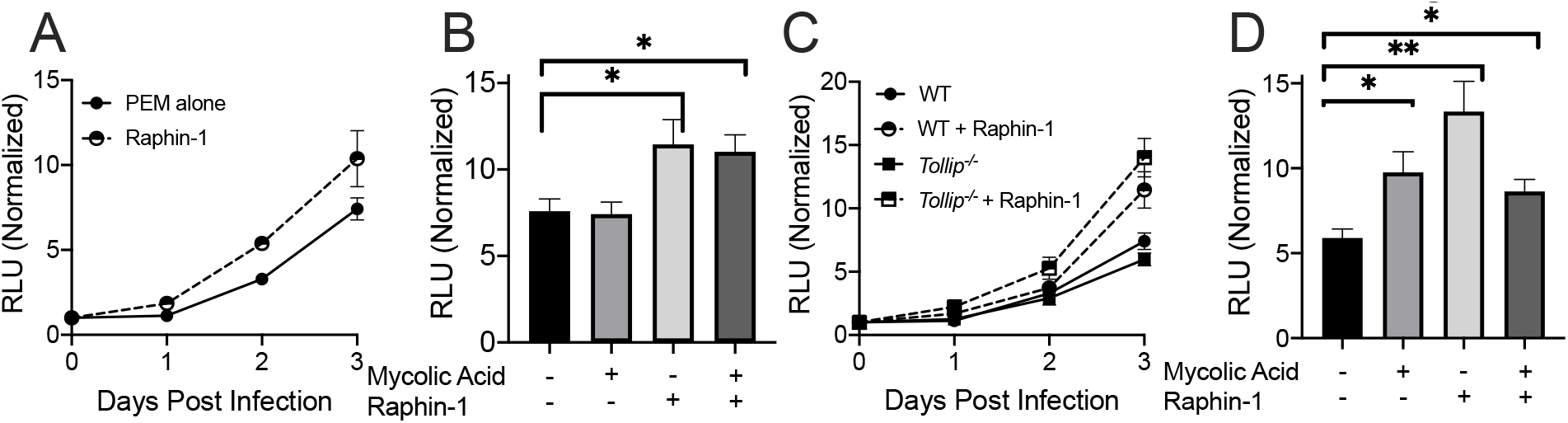
EIF2 Activation Impairs Mtb Control in Macrophages. Peritoneal extract macrophages (PEM) were plated and infected with Mtb expressing the lux luminescence plasmid (MOI 1). For each experiment luminescence was measured at the time points described. A) WT PEM were treated with vehicle control or raphin-1 (10μM) at the time of infection. Luminescence was measured over time. Statistical significance determined by 2-way ANOVA accounting for time and treatment. B) WT PEM were treated with mycolic acid (MA; 10μg/ml) for 72hr, then treated with either vector control or raphin-1 at the time of Mtb infection. Bars represent luminescence at 72hr after infection. Statistical significance determined by 2-way t-test. C) WT and *Tollip*^-/-^ PEM were treated with vehicle control or raphin-1, infected with Mtb, then luminescence was measured over time. Statistical significance determined by 3-way ANOVA accounting for TOLLIP expression, time, and treatment. D) *Tollip*^-/-^ PEM were treated with MA, raphin-1, or vehicle control as in B). Luminescence was measured 72h after infection. Statistical significance determined by two-sided unpaired t-test. Experiments were performed six times, all data is shown. * p < 0.05, ** p < 0.01.

### EIF2 Signaling Complex Proteins are Enriched in Human TB Granulomas

A defining characteristic of human granulomas is the presence of “caseum,” a lipid-rich center of necrosis (*37*). This central area of necrosis is surrounded by foam cells and other inflammatory cells induced by Mtb infection(*38*). Because we found that EIF2 signaling was induced by lipid accumulation and permissive to Mtb growth, we hypothesized that EIF2 complex genes are enriched in caseous granulomas. EIF2 signaling is activated by the phosphorylation of *EIF2S1*, at Ser48 and Ser51 by one of four upstream kinases which are responsive to different stress stimuli – *EIF2AK1* (HRI; heavy metals), *EIF2AK2* (PKR; dsRNA), *EIF2AK3* (PERK; unfolded protein/lipid), and *EIF2AK4* (GCN2; nutrient stress)(*13*). We evaluated the gene expression of *EIF2S1* and its four upstream kinases in human caseous granulomas (*16*). We compared mRNA transcript expression in lung tissues isolated from the caseous granulomas of individuals infected with TB with samples from healthy areas of the same lungs (n = 7 total). *EIF2AK1* (**Figure 8A**; FDR = 0.011), *EIF2AK2* (**Figure 8B**, FDR = 0.016), and *EIF2AK3* (**Figure 8C**, FDR = 0.002) demonstrated significantly increased expression in caseous granulomas. We did not detect significant increases in *EIF2S1* (**Figure 8D**). *EIF2AK4* and *TOLLIP* expression were also increased in TB granulomas (**Figure 8E-F**; FDR = 0.25 and 0.08, respectively), but did not meet our strict criteria for achieving statistical significance.

**Figure 8.**
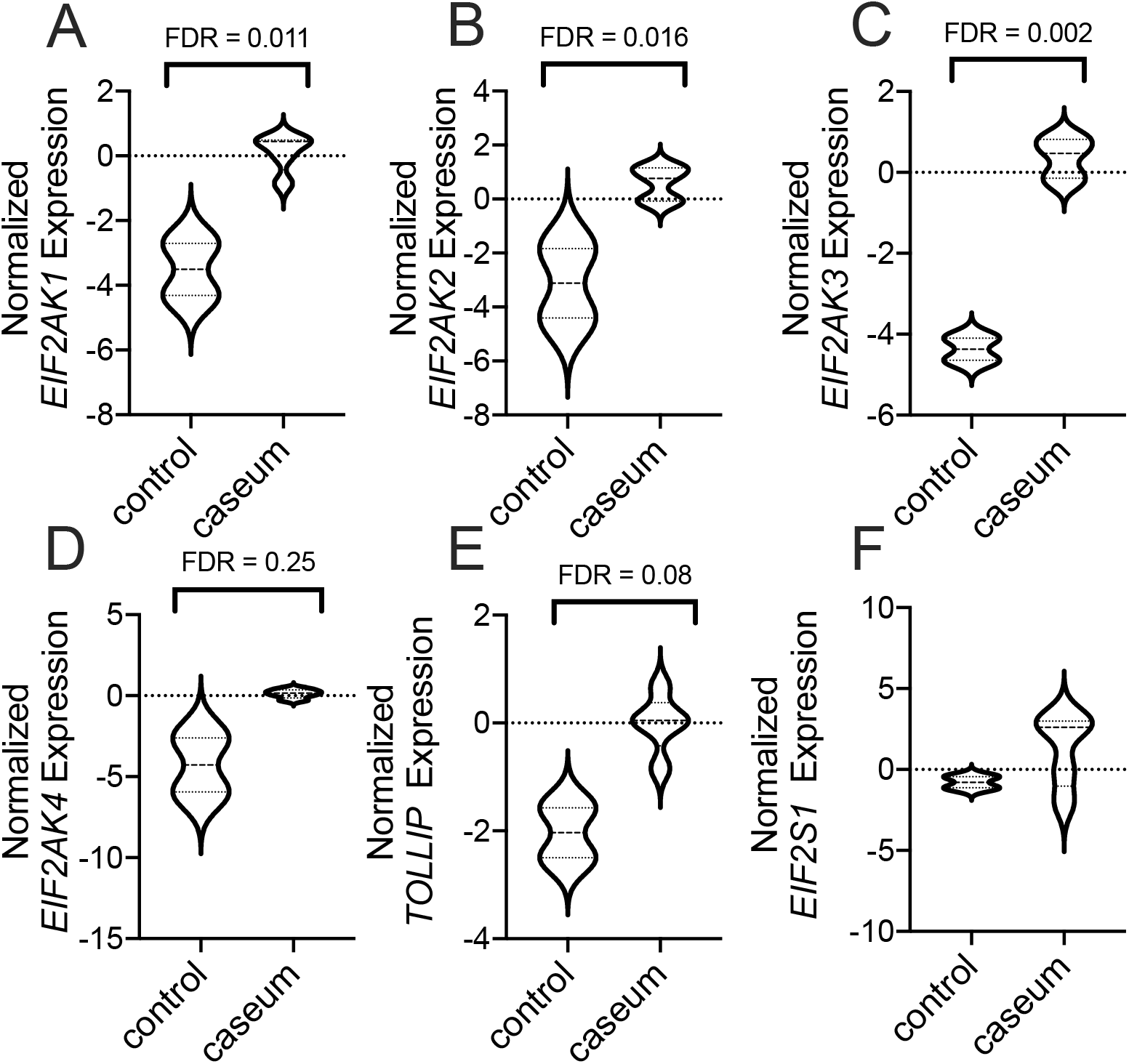
EIF2 Kinases are Enriched in Caseous TB Granulomas. Expression of A) *EIF2AK1*, B) *EIF2AK2*, C) *EIF2AK3*, D) *EIF2AK4*, E) *EIF2S1, and* F) *TOLLIP* in human caseous granuloma tissue, compared with healthy appearing lung tissue from different lobes from the same individuals. N = 7. Data are derived from the analysis of the publicly available GEO dataset GSE20050 using the indicated probe sets. FDR - false discovery rate. FDR is indicated above each data point.

## Discussion

Although intracellular lipid accumulation is a consistent feature of Mtb infection in humans, its relative absence in small animal models makes study of these phenotypes challenging. Mice lacking *Tollip* develop excessive lipid accumulation while maintaining increased innate immune activation. Therefore, this model of lipid accumulation is useful to evaluate excess lipid influences Mtb pathogenesis during prolonged infection and to define the molecular pathways responsible for these observed phenotypes. Computational analysis of lipid-loaded, Mtb-infected *Tollip*^-/-^ AM revealed that EIF2 signaling was the upstream pathway responsible for increased Mtb persistence, suggesting an important contribution of the ISR on Mtb pathogenesis *in vivo*. Importantly, lipids alone are sufficient to induce intracellular lipid accumulation and stress selectively in *Tollip*^-/-^ macrophages. Further, EIF2 kinases are enriched in human granulomas, providing human context for this observation.

EIF2 signaling prevents effective macrophage control of Mtb. ER stress complex proteins are increased in murine lungs after Mtb infection (*25*), but whether they are beneficial or harmful to the host has been challenging to determine previously (*16, 39*). Low level, acute EIF2 activity inhibits protein translation, except for a suite of proteins induced to maintain cellular survival, primarily through ATF4. ATF4 inhibits glycolysis and induces oxidative phosphorylation, (*40*) a metabolic pattern that is maladaptive in the Mtb-infected lung, and severe or prolonged EIF2 activation induces cellular death via CHOP (*41*). In a stressed state such as a caseous granuloma, all cells may die from stress in the necrotic center. Moreover, thresholds for cell death in response to stress are cell-specific, as secretory cells, such as plasma cells and islet cells, are more sensitive to stress (PMID 28951826;). Bringing these two observations together, the structure of human granulomas may be at least in part dictated by EIF2-induced stress responses, as they are classically lipid-rich, contain a central area of lipid necrosis, surrounded by foamy macrophages, then epithelioid macrophages and T and B cells. These data suggest a geographic and environmental gradient of ER stress signals contribute to necrosis, cellular distribution and morphology, and influence immune dysfunction and Mtb progression. Our study demonstrates that on balance, EIF2 signaling induced by lipid excess is maladaptive for Mtb control. Understanding how EIF2 signals impact granuloma formation may provide tools for improving the immune responses to Mtb.

How excess lipid induces stress to permit Mtb persistence within macrophages is not well described. In other cells, fatty acids induce cellular toxicity after destabilizing lysosomal membranes and increasing their permeability (*42*). Acid sphingomyelinase, a lysosomal enzyme, is required to cleave ceramide to sphingosine, which specifically induces lysosomal permeability (*43*). Permeability depletes calcium within the ER, which leads to mitochondrial oxidative stress (*44*). In settings of lysosomal permeability, cathepsins and other components of toxic granules may cross into the cytoplasm and induce macrophage necrosis and further exacerbate cellular stress. (*45, 46*). Recent zebrafish studies demonstrate that ER calcium flux and reactive oxygen species determine Mtb-induced macrophage cell death and Mtb escape from the macrophage (*21*). In Mtb-infected *Tollip*^-/-^ AM, we note strong enrichment of sphingolipid metabolic pathways. Combined with the observation that TOLLIP prevents lipid accumulation, we surmise that TOLLIP-deficient macrophages accumulate excess sphingosine to create lysosomal permeability and induce EIF2 signaling to permit Mtb replication. EIF2 then induces downstream cell cycle arrest and impaired metabolic adaptation, which provide a replicative niche for Mtb despite increasing inflammation in the surrounding tissues

In mixed bone marrow chimeras, *Tollip*^-/-^ AM developed increased Mtb burden 28 days post-infection. However, full knockout mice developed increased bacterial burden eight weeks post-infection. The reason for this observed difference is uncertain. Chimeric mice undergo full body radiation and bone marrow transplant, and this disruption of the bone marrow impacts future immune responses, including influencing the immune response to pathogens. Decreased TOLLIP expression is associated with impaired lung transplant engraftment, and so TOLLIP deficiency may selectively influence the immune response to mycobacteria after bone marrow reconstitution (*47*). However, we identified these phenotypes within multiple mouse models, and find this phenotype recapitulated in genetic studies of TOLLIP (*6, 28*). Alternately, intracellular carriage of mycobacteria within AM may more sensitively measure mycobacterial burden than total lung Mtb bacterial CFU, as intracellular replication within AM may precede replication in the lung overall. Thus, our observations in AMs using mixed bone marrow chimeric mice may represent a “leading indicator” toward the chronic, maladaptive phenotype. We focused on the impact of TOLLIP on macrophage biology in this study and discovered that it plays an AM-intrinsic role in lipid clearance, but TOLLIP is ubiquitously expressed and may influence the function of multiple immune cell subsets. Selective autophagy receptors alter dendritic cell function, particularly during EBV, influenza, yellow fever vaccine, and *L. monocytogenes* infection (*48–50*). ER stress response protein XBP1 prevents dendritic cell apoptosis and permits T cell activation during lipid excess in ovarian cancer models (*22*). TOLLIP may also impact T cell activation and differentiation. Another selective autophagy receptor, TAX1BP1, permits T cells to meet the energy demands of activation and proliferation (*51*), and autophagy supports regulatory T cell lineage stability by maintaining its metabolic homeostasis (*52*). Thus, TOLLIP may influence autophagy and cellular homeostasis, and lipids may impact cellular function, across cell types during Mtb infection.

Studies from the preantibiotic era regularly describe “post-primary” tuberculosis as an acute, paucibacillary, caseating lipoid pneumonia obstructing bronchioles in a “tree-in-bud” pattern (*53–55*), but its impact on Mtb pathogenesis is not clearly defined (*38*). In contrast, the classically defined cavitating granuloma is strongly associated with chronic established Mtb disease. Within this lipoid pneumonia, obstructed alveoli attract macrophages with a foamy appearance, leading to subsequent lipoid necrosis and granuloma formation around caseous necrosis. Pathologically, this obstructive lipoid pneumonia precedes the transition from asymptomatic to symptomatic TB disease (*38*). Thus, a combination of genetic factors that influence lipid clearance and physical factors that obstruct clearance of lipid debris may set the stage for progressive TB disease. Providing means to 1) promote ongoing lipid clearance or 2) relieve bronchial obstruction during the earliest phases of Mtb infection may prevent TB progression from early silent infection to symptomatic disease. Further, therapies to prevent this phenotype or resolve it may sensitize macrophages to immune signals, improving the effectiveness of vaccine candidates. Preventing lipid-induced EIF2 signaling via TOLLIP may be an effective host directed therapeutic strategy.

## Materials and Methods

### Reagents

A complete list of reagents can be found in **Table S1**.

### Mice

All mice were housed and maintained in specific pathogen-free conditions at the University of Washington and Seattle Children’s Research Institute, and all experiments were performed in compliance with the Institutional Animal Care and Use Committee from each institution. Mice used in the experiments were 6-12 weeks of age. B6.Cg-Tollip^tm1Kbns^/Cnrm (*Tollip*^-/-^) mice were obtained from the European Mutant Mouse Archive (www.infrafrontier.eu) (*56*). Mice were backcrossed 11 times on C57BL/6J background and were confirmed to be >99% C57BL/6J genetically by screening 150 SNP ancestry informative markers (Jax, Inc). Genotyping was performed using DNA primers for neomycin (Forward sequence: AGG ATC TCC TGT CAT CTC ACC TTG CTC CTG; Reverse sequence AAG AAC TCG TCA AGA AGG CGA TAG AAG GCG) and the first exon of TOLLIP (Forward sequence: AGC TAC TGG GAG GCC ATA CA; Reverse sequence: CGT GTA CGG GAG ACC CAT TT). Protein expression was confirmed in both knockout and backcrossed alleles by qPCR and Western blot. All wild type control mice were age-matched littermates to ensure a common genetic background.

### Model of Mtb aerosol infection

Aerosol infections were performed with wildtype H37Rv Mtb or H37Rv Mtb with an mCherry reporter plasmid. Mice were enclosed in a Glas-Col aerosol infection chamber and ~50-100 CFU were deposited into mouse lungs. Doses were confirmed using control mice by plating lung homogenates on 7H10 agar immediately after aersosol infection. Mice were sacrificed at indicated timepoint, and lungs were gently homogenized in PBS-containing 0.05%Tween using a gentleMacs dissociator (Miltenyi Biotec). Tissue homogenates were serially diluted on 7H10 agar and lung CFU was enumerated.

### Lung Cell Flow Cytometry

Lung single cell suspensions were washed and stained for viability with Zombie Aqua viability dye (BioLegend) for 10 min at room temperature in the dark. After incubation, 100 μl of a surface antibody cocktail diluted in 50% FACS buffer/50% 24G2 Fc block buffer was added and surface staining was performed for 30 min at 4°C. Antibody lists can be found in **Table S1**. The cells were washed once with FACS buffer and fixed with 2% paraformaldehyde for 1 h prior to analyzing on an LSRII flow cytometer (BD Biosciences). In some experiments, intracellular staining was performed after surface staining. Permeabilization was peformed with Fix-Perm buffer (eBiosciences) for minimum of 60 min before the addition of intracellular antibodies. Then cells were fixed with 2% paraformaldehyde for 1 h prior to analyzing on an LSRII flow cytometer (BD Biosciences). For cell sorting, mice were infected with Mtb H37Rv expressing mCherry (gift of David Sherman). 28 days after infection, mice were sacrificed, and AM were sorted on a BD FACSAria in a BSL-3 facility for infected and uninfected populations. The samples were spun and stored −80°C in Trizol.

### Chimera Generation

WT:*Tollip*^-/-^ mixed bone marrow chimeras were generated in the following manner: WT B6.SJL-Ptprca Pepcb/BoyJ (CD45.1+; Jax, Inc.) F1 mice were lethally irradiated (1000 cGy). A 1:1 mixture of CD3-depleted (Miltenyi Biotec) *Tollip*^-/-^ (CD45.2+) and F1 generation of C57BL/6J (CD45.1+45.2+) bone marrow was provided intravenously. For full bone marrow chimera generation, WT and *Tollip*^-/-^ mice were irradiated, followed by hemopoietic reconstitution by adoptive transfer of 5-10×10^6^ bone marrow cells via intravenous injection.

### Tissue Preparation and Evaluation

Mice were euthanized and lungs were gently homogenized in HEPES buffer containing Liberase Blendzyme 3 (70 μg/ml; Roche) and DNaseI (30 μg/ml; Sigma-Aldrich) using a gentleMacs dissociator (Miltenyi Biotec). The lungs were then incubated for 30 min at 37°C and then further homogenized a second time with the gentleMacs. The homogenates were filtered through a 70 μm cell strainer, pelleted for RBC lysis with RBC lysing buffer (Thermo), and resuspended in FACS buffer (PBS containing 2.5% FBS and 0.1% NaN_3_). To prepare organs for histology, lung sections were inflated to 15cm water pressure with 4% paraformaldehyde, fixed in the same solution, embedded in paraffin and 4□m sections were generated. Sections stained with hematoxylin and eosin were examined by a pathologist blinded to mouse genotype.

### Intracellular Mtb replication assay

Frozen Mtb was thawed and cultured on a shaking incubator for two doubling cycles. One day prior to infection, cultures were back-diluted into an optical density (OD) of 0.2–0.4 in 7H9 media supplemented with glycerol (4%), Middlebrook ADC Growth Supplement (100 mL/L), and Tween 80 (0.05%). At the time of infection, Mtb was filtered through a 5 μm syringe filter to remove bacterial clumps, and cells were inoculated at indicated MOI. The inoculum was prepared in RPMI-10 medium and applied to cells, which were centrifuged at 1200rpm for 5 minutes and incubated for 4 hours at 37°C. Supernatants were removed, and washed twice with prewarmed PBS (phosphate-buffered saline) to remove unbound bacteria, before adding per warmed RPMI supplemented with 10% FBS. Intracellular growth was determined using luminescence on a Synergy H4 multimode microplate reader (Biotek Instruments) daily from Day 0 to Day 3. In some experiments, 24hrs post infection cell culture supernatants were collected and filtered twice using 0.22μm filter and frozen at −80°C.

### Macrophage Preparation

Resident peritoneal cells, mainly consisting of *in vivo* differentiated macrophages, were isolated using standard methods (Zhang, X et al 2008). Briefly, 10ml of cold PBS was injected using 27g needle, and peritoneum was gently messaged. Cells were removed with PBS, centrifuged at 4° C and placed into warm RPMI media. PEMs (1×10^5^/well) were plated in 96-well plates in 200μL of antibiotic-free RPMI 1640 containing with 10% Fetal Bovine Serum (Atlas Biologics). Cell were rested for minimum 24hrs before further experimentation. Bone marrow was harvested from mice and grown in RPMI supplemented with 10% heat inactivated FBS and M-CSF (40ng/ml) in tissue culture treated plates. Cells were then incubated at 37° C for 6 to 7 days. Bone marrow-derived macrophages (BMDM) were used after 7 days of culture. BMDMs were detached by gently scrapping and cells were then plated in RPMI 1640 supplemented with 10% heat inactivated FBS.

### Cellular Studies

Cell-culture supernatants were collected at indicated time and frozen at −20 °C until analysis. The cytokine concentrations in the culture supernatant were determined using quantitative ELISA (Mouse TNF and IL-10 DuoSet; R&D Systems) as recommended by the manufacturer. For detection of lipid bodies, PEMs from WT and *Tollip*^-/-^ mice were plated (1×10^5^/well) in a glass bottom chamber slide. Lipid was added as previously described (*57*). Cells were washed twice and resuspended in PBS solution of HCS LipidTOX Deep Red Neutral Lipid stain (ThermoFisher Scientific) according to manufacturer instruction. After incubation, cells were washed twice with PBS and fixed in 2% PFA for 30min. Cells were mounted in medium containing DAPI stain (ProLong Gold, Thermo, Inc.). Lipid droplets were counted from 100 cells identified randomly selected high-powered fields.

### Western blotting

Immunoblots were performed as described previously (*58*). Briefly, 5-20μg of cell or tissue protein extract was separated by SDS-PAGE, transferred onto PVDF membranes and immunoblotted with primary antibodies, listed in **Table S1**. Secondary antibodies conjugated to horseradish peroxidase were added and luminescence was quantitated.

### Preparation of total RNA and sequencing

Total RNA from sorted cells was isolated using the manufacturer’s instructions (TRIzol, Invitrogen). The RNA purity and quantified was assessed by RNA-Tapestation (Agilent 4200). cDNA was prepared using SMART-Seq v4 Ultra Low Input RNA Kit (Takara Bio USA, Inc). RNA sequencing libraries were prepared using the Illumina TruSeq™ RNA Sample Preparation Kit (Illumina, San Diego, CA, USA). Samples were sequenced as 50 bp paired-end reads on a Illumina HiSeq 2500 and assessed using FastQC v0.11.8 to visualize sequence quality (*59*). Sequences were filtered using AdapterRemoval v2.3.1 to remove adapters, remove sequences with >1 ambiguous base, and trim ends to a 30+ Phred score (*60*). Sequences were aligned to the mouse genome mm10 using STAR v2.7.4a (*61*), and alignments were assessed with Picard v2.18.7 (*62*) and samtools v1.10 (*63*). Total reads in gene exons were quantified using featureCounts v2.0.1 (*64*).

### Differential expression analysis

Gene expression analyses were performed in R v3.6.1 with the tidyverse v1.3.0 (*65*). Overall, samples were very high-quality with > 32 million aligned sequences per sample and low variability in gene coverage (median coefficient of variation coverage < 0.6). Genes were filtered to protein coding genes with biomaRt (N = 21850) with > 0.1 counts per million (CPM) in at least three samples (N = 14215) (*66*). Counts were normalized for RNA composition using edgeR and log2 CPM voom normalized with quality weights using limma (*67, 68*). Differentially expressed genes were determined using limma using a contrasts model comparing WT and *Tollip*^-/-^ cells in uninfected and infected groups (**Table S2**). Genes with FDR < 0.3 for at least one contrast (N = 3899) were clustered into modules using WCGNA (*69*) with a minimum R-squared of 0.8 and minimum module size of 50. This resulted in 17 modules, leaving out 282 genes not grouped into any module (**Table S3**). Module expression was calculated as the mean expression of the log2 values of the genes within each module and assessed for differential expression using the same contrasts model in limma. Modules were functionally assessed using clusterProfiler v3.12.0 to compare them to the Broad Institute Hallmark gene sets in msigbr v7.0.1 by hypergeometric distribution (*70, 71*). The Benjamini–Hochberg correction for multiple comparisons was applied to P-values and significance was assessed at FDR < 0.05 for all tests.

### Statistics

For mouse experiments, groups of five mice were compared with one another unless otherwise indicated. Cellular measurements were compared using a two-sided Students’ *t* test unless otherwise specified. A value of *p* < 0.05 was considered a statistically significant result unless otherwise specified. Statistics were calculated using Prism version 8.0 (GraphPad, Inc.).

## Abbreviations

AM: alveolar macrophage
BafA: bafilomycin A
CFU: colony forming units
DC: dendritic cells
DEG: differentially-expressed genes
EIF2: eukaryotic initiation factor 2
ER: endoplasmic reticulum
IM: interstitial macrophages
LD: lipid droplet
MA: Mycolic acid
MDM: monocyte-derived macrophages
Mtb: *Mycobacterium tuberculosis*
PEM: peritoneal extract macrophages
PMN: neutrophils
TB: tuberculosis
TOLLIP: Toll-Interacting Protein
WT: wild-type
WGCNA: weighted gene coexpression network analysis

## Acknowledgements

The authors wish to thank the University of Washington Center for Lung Biology Histology and Imaging Core for their helpful advice on pathology staining and analysis. We are grateful to the Seattle Children’s Research Institute Cell Sorting Core for their assistance and technical support.

## Author Contributions

Conceptualization: SV, KU, SS JS; methodology: SV, SS, KU, MA, SD, JS; resources: JS, software: KDM, MA; validation: SV, JS, RE; formal analysis: MA, KDM; investigation: SV, SH, CP, GP, AL, AP, RP, JS; writing – original draft: SV, JS; writing – review and editing: KU, SS, MA, CP, JS; supervision: JS, KU, MA, SS; project administration: SV, JS; funding acquisition: JS, KU, SS, MA. Competing Interests: The authors declare no competing interests. Data and materials availability: All data from this study are available by request to. Abbreviations and description of bioinformatics tools and all bioinformatic data and code can be found at https://github.com/altman-lab/JS20.01.

## Supplemental Materials

**Figure S1.** TOLLIP deficiency induces a proinflammatory cytokine bias in Mtb-infected macrophages. Peritoneal exudate macrophages (PEM) were isolated, plated in tissue culture, and stimulated with LPS (10 ng/ml), PAM3 (250 ng/ml), and Mtb whole cell lysate (1mcg/ml) overnight. Cell culture supernatants (mean ± SD) were obtained and **A)** TNF and **B)** IL-10 concentrations were measured by ELISA. PEM were infected with Mtb H37Rv (MOI 2.5) overnight, then cell culture supernatants were collected and **C)** TNF, **D)** IL-1 β and **E)** IL-10 were measured by ELISA (mean ± SD). Two-sided t-test was used to determine statistical significance between groups. N=3/group, experiment was performed three times independently.

**Figure S2.** Lists flow cytometry gating strategy for Figure 3.

**Figure S3.** Lists flow cytometry gating strategy for Figure 5.

**Figure S4.** Describes sorting strategy for Figure 6.

**Figure S5.** Describes colocalization of Mtb and LC3 with Mtb in control and TOLLIP-deficient THP-1 cells.

**Figure S6.** Demonstrates increase in cellular stress in lipid-enriched, Mtb infected, Tollip-/- macrophages.

**Table S1.** Key Resources Table.

**Data File S1.** List of differentially expressed genes (DEG) in Mtb-infected and Mtb-uninfected *Tollip*^-/-^ AM compared to WT AM from the same mice.

**Data File S2.** List of genes in each WGCNA module comparing sorted WT and *Tollip*^-/-^ AM.

**Data File S3.** List of GSEA hallmark pathways significantly enriched in each WGCNA module comparing sorted WT and *Tollip*^-/-^ AM.

## References and Notes

1. K. Burns et al., Tollip, a new component of the IL-1RI pathway, links IRAK to the IL-1 receptor. Nat Cell Biol 2, 346–351 (2000).

2. K. Lu, I. Psakhye, S. Jentsch, Autophagic clearance of polyQ proteins mediated by ubiquitin-Atg8 adaptors of the conserved CUET protein family. Cell 158, 549–563 (2014).

3. M. L. Jongsma et al., An ER-Associated Pathway Defines Endosomal Architecture for Controlled Cargo Transport. Cell 166, 152–166 (2016).

4. K. Chen, R. Yuan, Y. Zhang, S. Geng, L. Li, Tollip Deficiency Alters Atherosclerosis and Steatosis by Disrupting Lipophagy. J Am Heart Assoc 6, (2017).

5. A. C. Martini et al., Amyloid-beta impairs TOM1-mediated IL-1R1 signaling. Proc Natl Acad Sci U S A 116, 21198–21206 (2019).

6. J. A. Shah et al., Genetic Variation in Toll-Interacting Protein Is Associated With Leprosy Susceptibility and Cutaneous Expression of Interleukin 1 Receptor Antagonist. J Infect Dis 213, 1189–1197 (2016).

7. J. A. Shah et al., A Functional TOLLIP Variant is Associated with BCG-Specific Immune Responses and Tuberculosis. Am J Respir Crit Care Med, (2017).

8. M. Ouimet et al., Mycobacterium tuberculosis induces the miR-33 locus to reprogram autophagy and host lipid metabolism. Nat Immunol 17, 677–686 (2016).

9. M. G. Gutierrez et al., Autophagy is a defense mechanism inhibiting BCG and Mycobacterium tuberculosis survival in infected macrophages. Cell 119, 753–766 (2004).

10. Y. Matsuzawa-Ishimoto, S. Hwang, K. Cadwell, Autophagy and Inflammation. Annu Rev Immunol 36, 73–101 (2018).

11. J. M. Kimmey et al., Unique role for ATG5 in neutrophil-mediated immunopathology during M. tuberculosis infection. Nature 528, 565–569 (2015).

12. B. Levine, N. Mizushima, H. W. Virgin, Autophagy in immunity and inflammation. Nature 469, 323–335 (2011).

13. J. Grootjans, A. Kaser, R. J. Kaufman, R. S. Blumberg, The unfolded protein response in immunity and inflammation. Nat Rev Immunol 16, 469–484 (2016).

14. M. Costa-Mattioli, P. Walter, The integrated stress response: From mechanism to disease. Science 368, (2020).

15. S. E. Bettigole, L. H. Glimcher, Endoplasmic reticulum stress in immunity. Annu Rev Immunol 33, 107–138 (2015).

16. M. J. Kim et al., Caseation of human tuberculosis granulomas correlates with elevated host lipid metabolism. EMBO Mol Med 2, 258–274 (2010).

17. B. Carow et al., Spatial and temporal localization of immune transcripts defines hallmarks and diversity in the tuberculosis granuloma. Nat Commun 10, 1823 (2019).

18. C. Koumenis et al., Regulation of protein synthesis by hypoxia via activation of the endoplasmic reticulum kinase PERK and phosphorylation of the translation initiation factor eIF2alpha. Mol Cell Biol 22, 7405–7416 (2002).

19. N. Mizushima, M. Komatsu, Autophagy: renovation of cells and tissues. Cell 147, 728–741 (2011).

20. D. G. Russell, L. Huang, B. C. VanderVen, Immunometabolism at the interface between macrophages and pathogens. Nat Rev Immunol 19, 291–304 (2019).

21. F. J. Roca, L. J. Whitworth, S. Redmond, A. A. Jones, L. Ramakrishnan, TNF Induces Pathogenic Programmed Macrophage Necrosis in Tuberculosis through a Mitochondrial-Lysosomal-Endoplasmic Reticulum Circuit. Cell 178, 1344–1361 e1311 (2019).

22. A. Kaser et al., XBP1 links ER stress to intestinal inflammation and confers genetic risk for human inflammatory bowel disease. Cell 134, 743–756 (2008).

23. J. Oh et al., Endoplasmic reticulum stress controls M2 macrophage differentiation and foam cell formation. J Biol Chem 287, 11629–11641 (2012).

24. A. J. Olive, C. M. Sassetti, Metabolic crosstalk between host and pathogen: sensing, adapting and competing. Nat Rev Microbiol 14, 221–234 (2016).

25. T. A. Seimon et al., Induction of ER stress in macrophages of tuberculosis granulomas. PLoS One 5, e12772 (2010).

26. Y. J. Lim et al., Mycobacterium tuberculosis 38-kDa antigen induces endoplasmic reticulum stress-mediated apoptosis via toll-like receptor 2/4. Apoptosis 20, 358–370 (2015).

27. S. Liang et al., BAG2 ameliorates endoplasmic reticulum stress-induced cell apoptosis in Mycobacterium tuberculosis-infected macrophages through selective autophagy. Autophagy 16, 1453–1467 (2020).

28. J. A. Shah et al., Human TOLLIP Regulates TLR2 and TLR4 Signaling and Its Polymorphisms Are Associated with Susceptibility to Tuberculosis. Journal of immunology 189, 1737–1746 (2012).

29. G. T. Consortium, Human genomics. The Genotype-Tissue Expression (GTEx) pilot analysis: multitissue gene regulation in humans. Science 348, 648–660 (2015).

30. J. A. Shah et al., TOLLIP deficiency is associated with increased resistance to Legionella pneumophila pneumonia. Mucosal Immunol 12, 1382–1390 (2019).

31. M. Guilliams et al., Alveolar macrophages develop from fetal monocytes that differentiate into long-lived cells in the first week of life via GM-CSF. J Exp Med 210, 1977–1992 (2013).

32. S. B. Cohen et al., Alveolar Macrophages Provide an Early Mycobacterium tuberculosis Niche and Initiate Dissemination. Cell Host Microbe 24, 439–446 e434 (2018).

33. A. Liberzon et al., The Molecular Signatures Database (MSigDB) hallmark gene set collection. Cell Syst 1, 417–425 (2015).

34. M. Ouimet et al., Autophagy regulates cholesterol efflux from macrophage foam cells via lysosomal acid lipase. Cell Metab 13, 655–667 (2011).

35. D. Scheuner et al., Translational control is required for the unfolded protein response and in vivo glucose homeostasis. Mol Cell 7, 1165–1176 (2001).

36. A. Krzyzosiak et al., Target-Based Discovery of an Inhibitor of the Regulatory Phosphatase PPP1R15B. Cell 174, 1216–1228 e1219 (2018).

37. R. L. Hunter, On the pathogenesis of post primary tuberculosis: the role of bronchial obstruction in the pathogenesis of cavities. Tuberculosis (Edinb) 91 Suppl 1, S6–10 (2011).

38. R. L. Hunter, Tuberculosis as a three-act play: A new paradigm for the pathogenesis of pulmonary tuberculosis. Tuberculosis (Edinb) 97, 8–17 (2016).

39. D. G. Russell, P. J. Cardona, M. J. Kim, S. Allain, F. Altare, Foamy macrophages and the progression of the human tuberculosis granuloma. Nat Immunol 10, 943–948 (2009).

40. E. Balsa et al., ER and Nutrient Stress Promote Assembly of Respiratory Chain Supercomplexes through the PERK-eIF2alpha Axis. Mol Cell 74, 877–890 e876 (2019).

41. J. Han et al., ER-stress-induced transcriptional regulation increases protein synthesis leading to cell death. Nat Cell Biol 15, 481–490 (2013).

42. F. G. Almaguel et al., Lipotoxicity-mediated cell dysfunction and death involve lysosomal membrane permeabilization and cathepsin L activity. Brain Res 1318, 133–143 (2010).

43. F. J. Roca, L. Ramakrishnan, TNF dually mediates resistance and susceptibility to mycobacteria via mitochondrial reactive oxygen species. Cell 153, 521–534 (2013).

44. S. Xu et al., Palmitate induces ER calcium depletion and apoptosis in mouse podocytes subsequent to mitochondrial oxidative stress. Cell Death Dis 6, e1976 (2015).

45. K. F. Ferri, G. Kroemer, Organelle-specific initiation of cell death pathways. Nat Cell Biol 3, E255–263 (2001).

46. K. Kagedal, U. Johansson, K. Ollinger, The lysosomal protease cathepsin D mediates apoptosis induced by oxidative stress. FASEB J 15, 1592–1594 (2001).

47. E. Cantu et al., Protein Quantitative Trait Loci Analysis Identifies Genetic Variation in the Innate Immune Regulator TOLLIP in Post-Lung Transplant Primary Graft Dysfunction Risk. Am J Transplant 16, 833–840 (2016).

48. R. Ravindran et al., Vaccine activation of the nutrient sensor GCN2 in dendritic cells enhances antigen presentation. Science 343, 313–317 (2014).

49. D. Schmid, M. Pypaert, C. Munz, Antigen-loading compartments for major histocompatibility complex class II molecules continuously receive input from autophagosomes. Immunity 26, 79–92 (2007).

50. H. K. Lee et al., In vivo requirement for Atg5 in antigen presentation by dendritic cells. Immunity 32, 227–239 (2010).

51. M. I. Whang et al., The Ubiquitin Binding Protein TAX1BP1 Mediates Autophagasome Induction and the Metabolic Transition of Activated T Cells. Immunity 46, 405–420 (2017).

52. J. Wei et al., Autophagy enforces functional integrity of regulatory T cells by coupling environmental cues and metabolic homeostasis. Nat Immunol 17, 277–285 (2016).

53. R. L. Hunter, Pathology of post primary tuberculosis of the lung: an illustrated critical review. Tuberculosis (Edinb) 91, 497–509 (2011).

54. V. Cornil, L. Ranvier, A manual of pathological histology translated with notes and additions by EO Shakespeare and JHC Simms. Philadephia: Henry C Lea, 394–445 (1880).

55. J. G. Im, H. Itoh, K. S. Lee, M. C. Han, CT-pathology correlation of pulmonary tuberculosis. Crit Rev Diagn Imaging 36, 227–285 (1995).

56. A. Didierlaurent et al., Tollip regulates proinflammatory responses to interleukin-1 and lipopolysaccharide. Mol Cell Biol 26, 735–742 (2006).

57. P. Peyron et al., Foamy macrophages from tuberculous patients’ granulomas constitute a nutrient-rich reservoir for M. tuberculosis persistence. PLoS Pathog 4, e1000204 (2008).

58. S. A. Soleimanpour et al., The diabetes susceptibility gene Clec16a regulates mitophagy. Cell 157, 1577–1590 (2014).

59. S. Andrews. (2010).

60. M. Schubert, S. Lindgreen, L. Orlando, AdapterRemoval v2: rapid adapter trimming, identification, and read merging. BMC research notes 9, 88–88 (2016).

61. A. Dobin et al., STAR: ultrafast universal RNA-seq aligner. Bioinformatics (Oxford, England) 29, 15–21 (2013).

62. (Broad Institute, 2019).

63. H. Li et al., The Sequence Alignment/Map format and SAMtools. Bioinformatics (Oxford, England) 25, 2078–2079 (2009).

64. Y. Liao, G. K. Smyth, W. Shi, featureCounts: an efficient general purpose program for assigning sequence reads to genomic features. Bioinformatics 30, 923–930 (2013).

65. H. Wickham et al., Welcome to the Tidyverse. Journal of Open Source Software 4, 1686–1686 (2019).

66. S. Durinck et al., BioMart and Bioconductor: a powerful link between biological databases and microarray data analysis. Bioinformatics 21, 3439–3440 (2005).

67. M. D. Robinson, D. J. McCarthy, G. K. Smyth, edgeR: a Bioconductor package for differential expression analysis of digital gene expression data. Bioinformatics (Oxford, England) 26, 139–140 (2010).

68. M. E. Ritchie et al., limma powers differential expression analyses for RNA-sequencing and microarray studies. Nucleic acids research 43, e47–e47 (2015).

69. P. Langfelder, S. Horvath, WGCNA: an R package for weighted correlation network analysis. BMC Bioinformatics 9, 559 (2008).

70. G. Yu, L.-G. Wang, Y. Han, Q.-Y. He, clusterProfiler: an R package for comparing biological themes among gene clusters. Omics: a journal of integrative biology 16, 284–287 (2012).

71. I. Dolgalev. (2019).

