## Supplementary figures and images for "TOLLIP resolves lipid-induced EIF2 signaling in alveolar macrophages for durable *Mycobacterium tuberculosis* protection"

### Supplemental Figure 1

#### Supplemental Figure 1

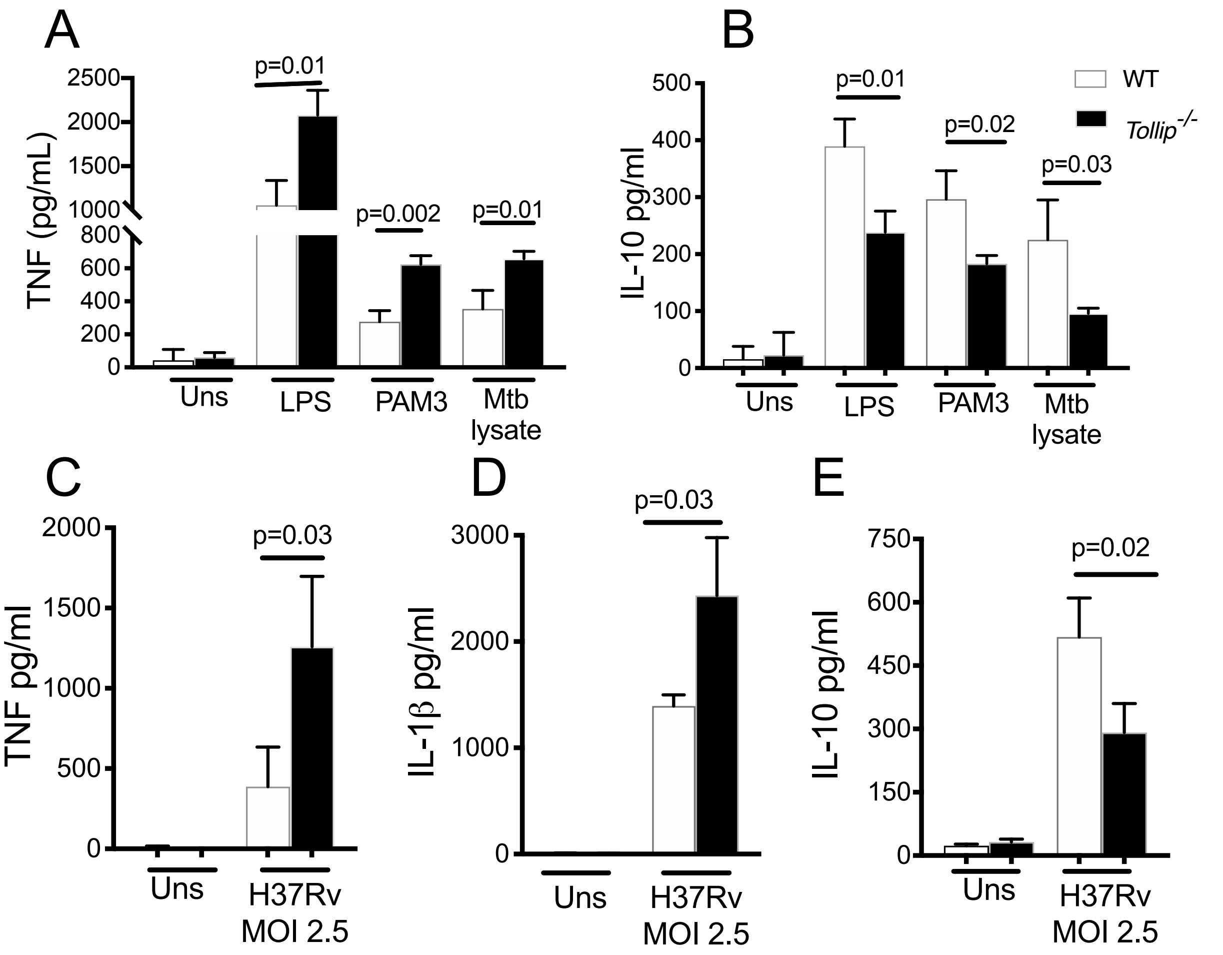

### Supplemental Figure 4

#### Supplemental Figure 4

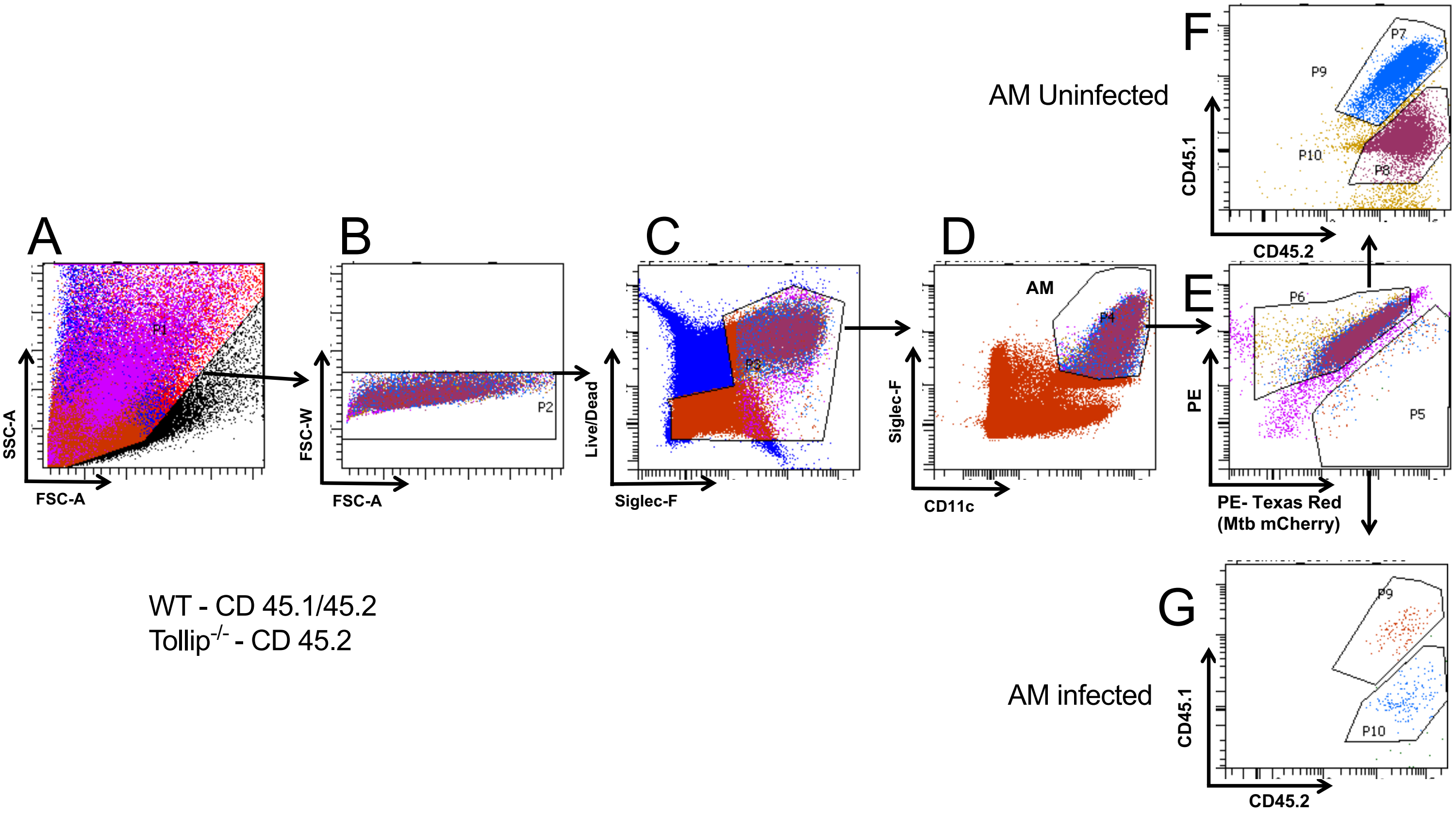

### Supplemental Figure 5

#### Supplemental Figure 5

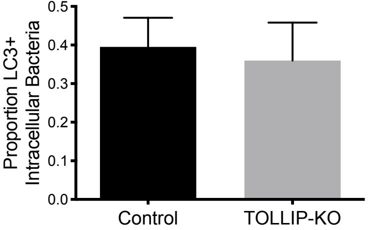


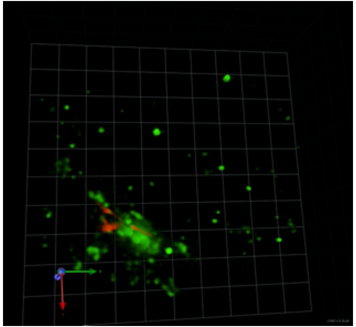

Control

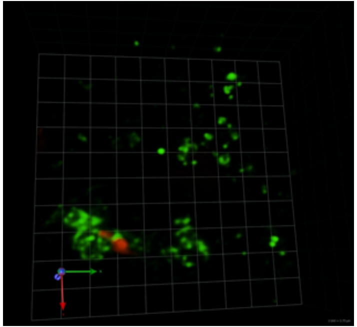

TOLLIP-KO

### Supplemental Figure 6

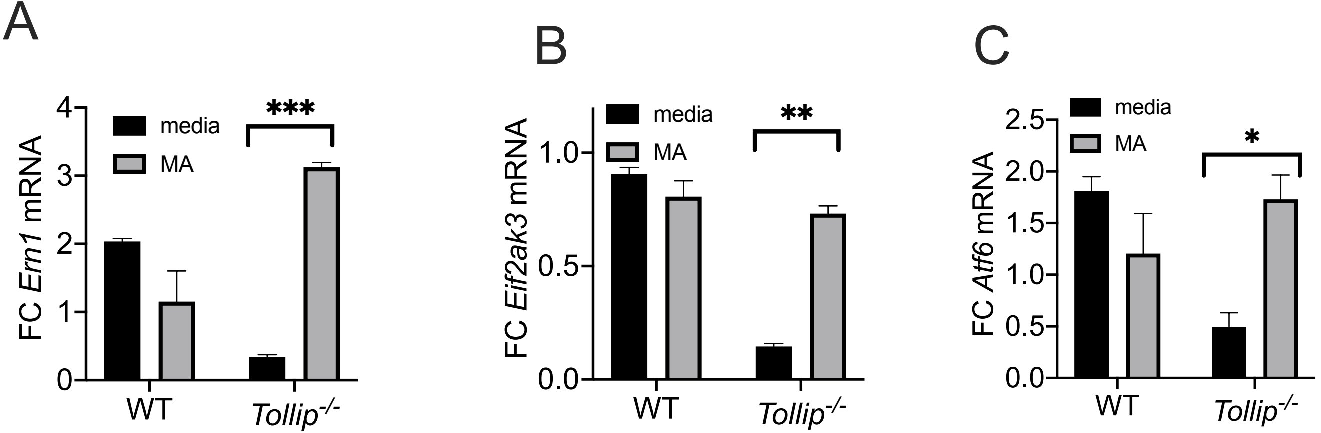
