## Supplemental Figure 3 for "TOLLIP resolves lipid-induced EIF2 signaling in alveolar macrophages for durable *Mycobacterium tuberculosis* protection"

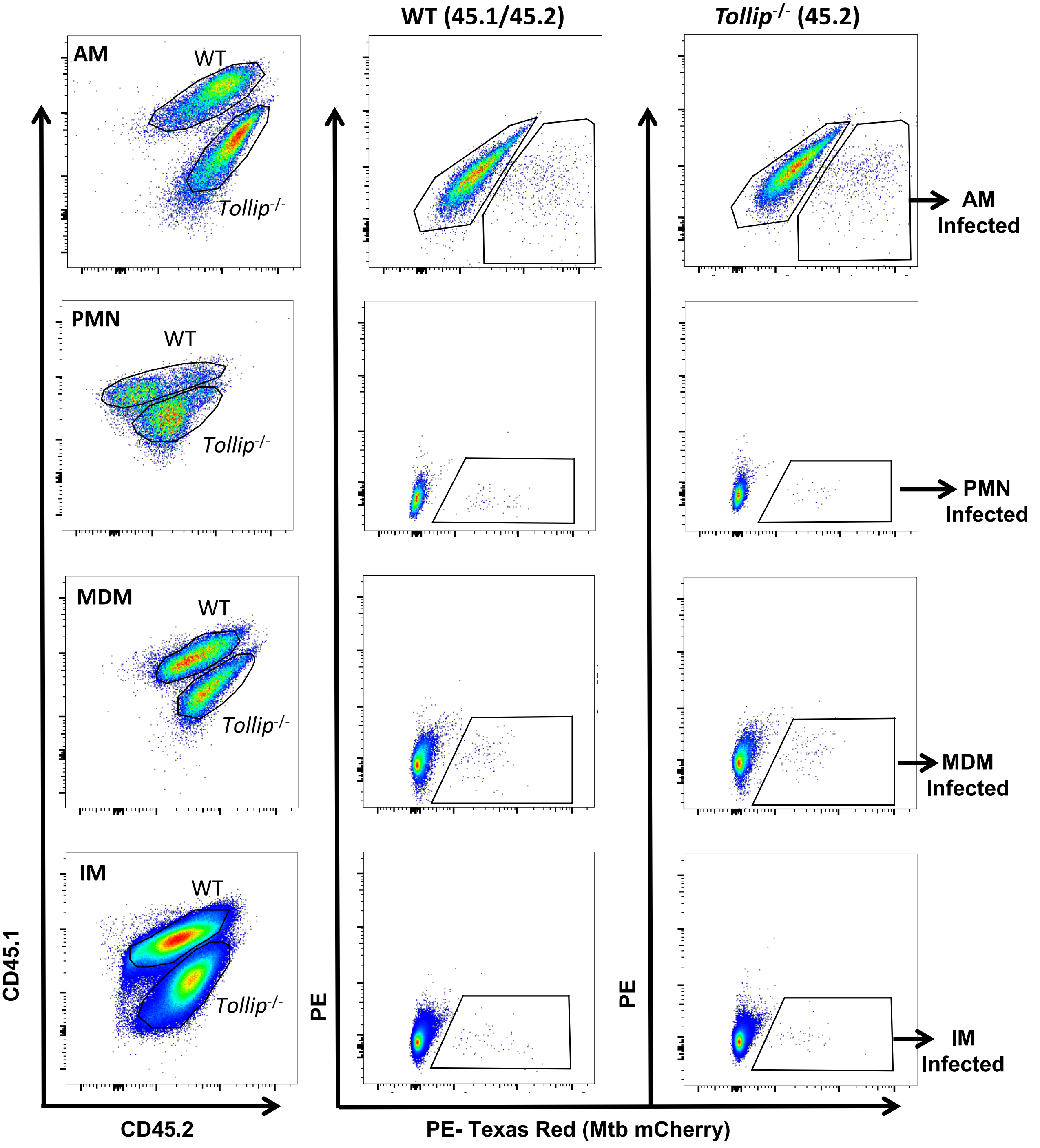


AM: Alveolar macrophage, PMN: Neutrophil, MDM: Monocyte-derived macrophages, IM: Interstitial macrophage.
