## Supplemental Materials for "TOLLIP resolves lipid-induced EIF2 signaling in alveolar macrophages for durable *Mycobacterium tuberculosis* protection"

**Figure S1.** TOLLIP deficiency induces a proinflammatory cytokine bias in Mtb-infected macrophages. Peritoneal exudate macrophages (PEM) were isolated, plated in tissue culture, and stimulated with LPS (10 ng/ml), PAM3 (250 ng/ml), and Mtb whole cell lysate (1mcg/ml) overnight. Cell culture supernatants (mean ± SD) were obtained and **A)** TNF and **B)** IL-10 concentrations were measured by ELISA. PEM were infected with Mtb H37Rv (MOI 2.5) overnight, then cell culture supernatants were collected and **C)** TNF, **D)** IL-1$\beta$ and **E)** IL-10 were measured by ELISA (mean ± SD). Two-sided t-test was used to determine statistical significance between groups. N=3/group, experiment was performed three times independently.

**Figure S2.** Lists flow cytometry gating strategy for Figure 3.

**Figure S3.** Lists flow cytometry gating strategy for Figure 5.

**Figure S4.** Describes sorting strategy for Figure 6.

**Figure S5.** Describes colocalization of Mtb and LC3 with Mtb in control and TOLLIP-deficient THP-1 cells.

**Figure S6.** Demonstrates increase in cellular stress in lipid-enriched, Mtb infected, Tollip-/- macrophages.

**Data File S1.** List of differentially expressed genes (DEG) in Mtb- infected and Mtb-uninfected *Tollip*$^{-/-}$ AM compared to WT AM from the same mice.

**Data File S2.** List of genes in each WGCNA module comparing sorted WT and *Tollip*$^{-/-}$ AM.

**Data File S3.** List of GSEA hallmark pathways significantly enriched in each WGCNA module comparing sorted WT and *Tollip*$^{-/-}$ AM.

**Table S1. Key Resources.**

| REAGENT or RESOURCE | SOURCE | IDENTIFIER |
|---|---|---|
| Antibodies | | |
| CD80 | Invitrogen | 11-0801-85 |
| CD40 | BioLegend | 124624 |
| CD11b | BioLegend | 101216 |
| SiglecF | BD | 562681 |
| Viability – Zombie Aqua | BioLegend | 423102 |
| Ly6G | BioLegend | 127614 |
| MHCII (I-A/I-E Antibody) | BioLegend | 107622 |
| NOS2 | Invitrogen | 47-5920-82 |
| CD11c | BioLegend | 117339 |
| CD103 | BD | 564322 |
| CD45.2 | BioLegend | 109828 |
| MHCII | BioLegend | 107620 |
| Ly6G | BD | 561105 |
| CD45.1 | Invitrogen | A14733 |
| CD45.2 | BioLegend | 109814 |
| CD11b | BioLegend | 101243 |
| CD45.1 | Invitrogen | 47-0453-82 |
| Bacterial and Virus Strains | | |
| M. tuberculosis H37Rv strain | David Sherman | |
| M. tuberculosis H37Rv strain – mCherry reporter | David Sherman | |
| M. tuberculosis H37Rv strain – *lux* expression reporter | Jeffrey Cox | |
| Critical Commercial Assays | | |
| Mouse TNF Duoset | R&D Systems | |
| Mouse IL-10 Duoset | R&D Systems | |

| | | |
|---|---|---|
| LipidTox Deep Red Neutral Lipid Stain | ThermoFisher | H34477 |
| Deposited Data | | |
| RNA-seq code | Shah lab | https://github.com/altman-lab/JS20.01 |
| Experimental Models: Cell Lines | | |
| THP-1, empty vector control | Shah lab | |
| THP-1, TOLLIP-knockout | Shah lab | |
| Experimental Models: Organisms/Strains | | |
| B6.Cg-Tollip$^{tm1Kbns}$/Cnrm | European Mutant Mouse Archive (www.infrafrontier.eu) | |
| B6.SJL-Ptprca Pepcb/BoyJ | Jax, Inc. | Stock # 002014 |
| Oligonucleotides | | |
| Neomycin for – mouse genotyping AGG ATC TCC TGT CAT CTC ACC TTG CTC CTG | IDT | |
| Neomycin rev – mouse genotyping AAG AAC TCG TCA AGA AGG CGA TAG AAG GCG | IDT | |
| Software and Algorithms | | |
| FastQC v.0.11.8 | [1] | |
| AdapterRemoval v2.3.1 | [2] | |
| STAR v2.7.4 | [3] | |
| Picard v2.18.7 | [4] | |
| Samtools v1.10 | [5] | |
| featureCounts v2.0.1 | [6] | |
| R v3.6.1 | | |
| Tidyverse v1.3.0 | [7] | |
| biomaRt | [8] | |
| limma | [9,10]. | |
| edgeR | [9,10]. | |
| WGCNA | [11] | |
| Clusterprofiler v3.12.0 | [12] | |
| Misbr v7.0.1 | [13] | |

1. Andrews S. FastQC: A Quality Control Tool for High Throughput Sequence Data. 2010.
2. Schubert M, Lindgreen S, Orlando L. AdapterRemoval v2: rapid adapter trimming, identification, and read merging. BMC research notes 2016;9:88-.
3. Dobin A, Davis CA, Schlesinger F, et al. STAR: ultrafast universal RNA-seq aligner. Bioinformatics (Oxford, England) 2013;29:15-21.
4. Picard Toolkit. Broad Institute; 2019.
5. Li H, Handsaker B, Wysoker A, et al. The Sequence Alignment/Map format and SAMtools. Bioinformatics (Oxford, England) 2009;25:2078-9.

6. Liao Y, Smyth GK, Shi W. featureCounts: an efficient general purpose program for assigning sequence reads to genomic features. Bioinformatics 2013;30:923-30.
7. Wickham H, Averick M, Bryan J, et al. Welcome to the Tidyverse. Journal of Open Source Software 2019;4:1686-.
8. Durinck S, Moreau Y, Kasprzyk A, et al. BioMart and Bioconductor: a powerful link between biological databases and microarray data analysis. Bioinformatics 2005;21:3439-40.
9. Robinson MD, McCarthy DJ, Smyth GK. edgeR: a Bioconductor package for differential expression analysis of digital gene expression data. Bioinformatics (Oxford, England) 2010;26:139-40.
10. Ritchie ME, Phipson B, Wu D, et al. limma powers differential expression analyses for RNA-sequencing and microarray studies. Nucleic acids research 2015;43:e47-e.
11. Langfelder P, Horvath S. WGCNA: an R package for weighted correlation network analysis. BMC Bioinformatics 2008;9:559.
12. Yu G, Wang L-G, Han Y, He Q-Y. clusterProfiler: an R package for comparing biological themes among gene clusters. Omics : a journal of integrative biology 2012;16:284-7.
13. Dolgalev I. msigdbr: MSigDB Gene Sets for Multiple Organisms in a Tidy Data Format. 2019.
